# Maternal humoral factors modulate offspring gut immune homeostasis to mitigate diabetes development

**DOI:** 10.1101/2024.08.28.604371

**Authors:** Erin Strachan, Luisa Pessoa-Soares, Tingting Ju, Alejandro Schcolnik-Cabrera, Henry Wang, Masoud Akbari, Paulo Jose Basso, Daniel A. Winer, Carol Huang, Olivier Julien, Benjamin P. Willing, Xavier Clemente-Casares, Sue Tsai

## Abstract

Environmental risk factors possess the potential to modulate the pathogenesis of type I diabetes (T1D). Foremost among these factors are early life influences impacting the gastrointestinal (GI) tract. During infancy, both the microbiota and immune system are influenced by maternal factors contributing to key events in the neonatal GI tract. Despite the well-known importance of maternal factors on infant immune development, whether maternal immune dysregulation and dysbiosis can perpetuate the same in offspring remains largely unknown. To explore how these maternal factors impact offspring disease development, we used IgA-deficiency induced maternal dysbiosis in Non-Obese Diabetic (NOD) dams to study T1D development in their progeny. We found that maternal dysbiosis and absence of IgA led to changes in IgA-sufficient offspring immune development resulting in heightened GI immune activity. Maternal dysbiosis also contributed to altered microbiome establishment in progeny, such that pups exhibited reduced colonic abundance of *Akkermansia muciniphila* and *Clostridoides difficile*. In adulthood, these mice exhibited a lowered incidence of T1D. This protection was replicated by fostering high incidence offspring to dysbiotic dams, prompting us to propose that altered breast milk composition in dysbiotic dams can influence immune development and microbiome establishment in offspring, contributing to T1D resistance.

## Introduction

The immune system is tasked with identifying potentially harmful foreign entities while perpetuating tolerance to self tissues. Proper immune development, education and regulation are key to the successful maintenance of this balance, and errors in these processes can give rise to a myriad of autoimmune diseases such as type 1 diabetes (T1D). Defective central and peripheral tolerance mechanisms are at the core of T1D pathology, culminating in the inappropriate activation of self-reactive lymphocytes targeting the beta cells of the pancreas^1^. T1D is estimated to affect 8.42 million people worldwide, disproportionately affecting younger individuals^2^. Susceptibility to the disease is complex, involving a consortium of genetic, environmental, dietary and lifestyle determinants that can culminate in either resistance to or heightened risk of T1D development^3,4^. T1D is somewhat unique amongst autoimmune diseases in that by the time the patient receives a diagnosis, the affected tissue has already been irreparably damaged. Thus, there have been considerable efforts aimed at understanding the early events leading to disease onset, with the hope of establishing early intervention strategies to prevent beta cell loss in individuals at risk^5^. In parallel, there has been increasing pressure to elucidate key non-genetic factors that influence disease susceptibility, as recent decades have seen a significant rise in the proportion of newly diagnosed individuals lacking genetic loci traditionally linked with high T1D risk^6^.

One particularly intriguing non-genetic risk factor that has come to light through familial T1D studies may hold key insights to the establishment of early intervention strategies. The increased T1D risk associated with being born to an affected parent is reduced 4-fold when the affected parent is the mother vs the father^7–17^. This raises a multitude of questions regarding how non-genetic maternal factors influence disease susceptibility in offspring^18^. Maternal provisions are key contributors to the health and well-being of neonates. Breast milk immuno-modulatory factors play a large role in infant immune development and education^19–25^ and maternal microbes are primary colonizers of the early microbiome in infants^26,27^. Moreover, one’s microbiome plays a pivotal role in modulating immunity within the gut as well as throughout the body^21,22,28^. This is particularly true during the neonatal period, when microbiome establishment modulates immune education, development and regulation with lifelong impact^21,22,29^. The importance of establishing a healthy microbiome during early life is underscored by the multitude of studies linking early microbiome perturbations with immune-dysregulatory diseases contracted later in life^30^.

T1D susceptibility is strongly associated with changes in the gut microbiota^31^. Differences in microbial diversity, composition and activity are seen prior to diagnosis^32–36^ and can contribute to early disease processes^37,38^. Additionally, disease associated physiological changes in the gut environment^32,39–41^ may reciprocally favor a microbial community that promotes T1D. To date, investigation into the relationship between T1D-associated maternal dysbiosis and offspring disease development remains limited. The observation that diabetic mothers confer disease protection to their children prompted us to explore if and how disease-associated maternal factors modulate infant microbiome establishment, immune development and disease susceptibility, particularly within a genetically permissive setting of familial T1D.

Immunoglobulin A (IgA) constitutes a dominant mucosal antibody. It is constitutively secreted in large quantities in dimeric form by plasma cells (PC) into mucosal surfaces and plays a pivotal role in sustaining commensalism and antimicrobial defence^42–45^. We previously showed that genetic IgA deficiency in mice also resulted in a dysbiotic gut and exacerbated obesity-related inflammation and insulin resistance^46,47^. In autoimmune central nervous system (CNS) inflammation, IgA-producing PC are found in the CNS and play an important protective role^48,49^. Interestingly, selective IgA deficiency is associated with autoimmune diabetes and celiac disease^50–54^, and changes in IgA levels and function (i.e., binding quality) accompany T1D^55–59^. These changes can indirectly impact autoimmunity by modulating the production of immunomodulatory microbial metabolites in the gut and systemically^60^. Nonetheless, it is unclear whether disease-associated IgA alterations manifest as a cause or a consequence of dysbiosis in T1D.

Here, we set out to investigate the impact of maternal IgA deficiency and associated dysbiosis on early gastrointestinal (GI) environment, immune development and T1D progression in genetically-susceptible offspring. We introduced a genetic deletion of the IgA constant region into the non-obese diabetic (NOD) mouse model to induce microbiome perturbations, resulting in T1D-susceptible animals that are deficient in IgA. To our surprise, maternal IgA deficiency significantly reduced T1D development in the pups. Disease protection was linked to altered maternal humoral immunity and offspring microbiome changes, which are enacted during the postnatal early life period leading up to weaning. Our findings demonstrate the presence of maternal humoral factors that afford dominant protection against disease.

## Results

### Multi-generational IgA deficiency reduces microbiome diversity without altering disease incidence

We first assessed the bacterial communities in fecal material from 12 week old NOD.IgA KO female progeny of ≥3 generations of IgA deficient crosses compared to NOD controls via 16S rRNA sequencing. Comparison of the two cohorts, each consisting of mice from four different litters, revealed significant changes (FIGURE 1A), with the NOD.IgA KO fecal microbiome exhibiting reduced alpha diversity (FIGURE 1B). Depleted taxa included *Lachnospiraceae* (12 spp.), *Muribaculaceae* (12 spp.) *Alistipes* (2 spp.), and *Oscillospiraceae* (3 spp.) as well as the homeostasis-promoting species *Lactobacillus murinus*^61^ (FIGURE 1C). Conversely, the decrease in diversity in NOD.IgA KO was accompanied by the outgrowth of a few taxa, including one species each from the family *Lachnospiraceae*, the genus *Bacteroides*, and the enteric pathogen *Clostridioides difficile* (FIGURE 1C).

**FIGURE 1.**
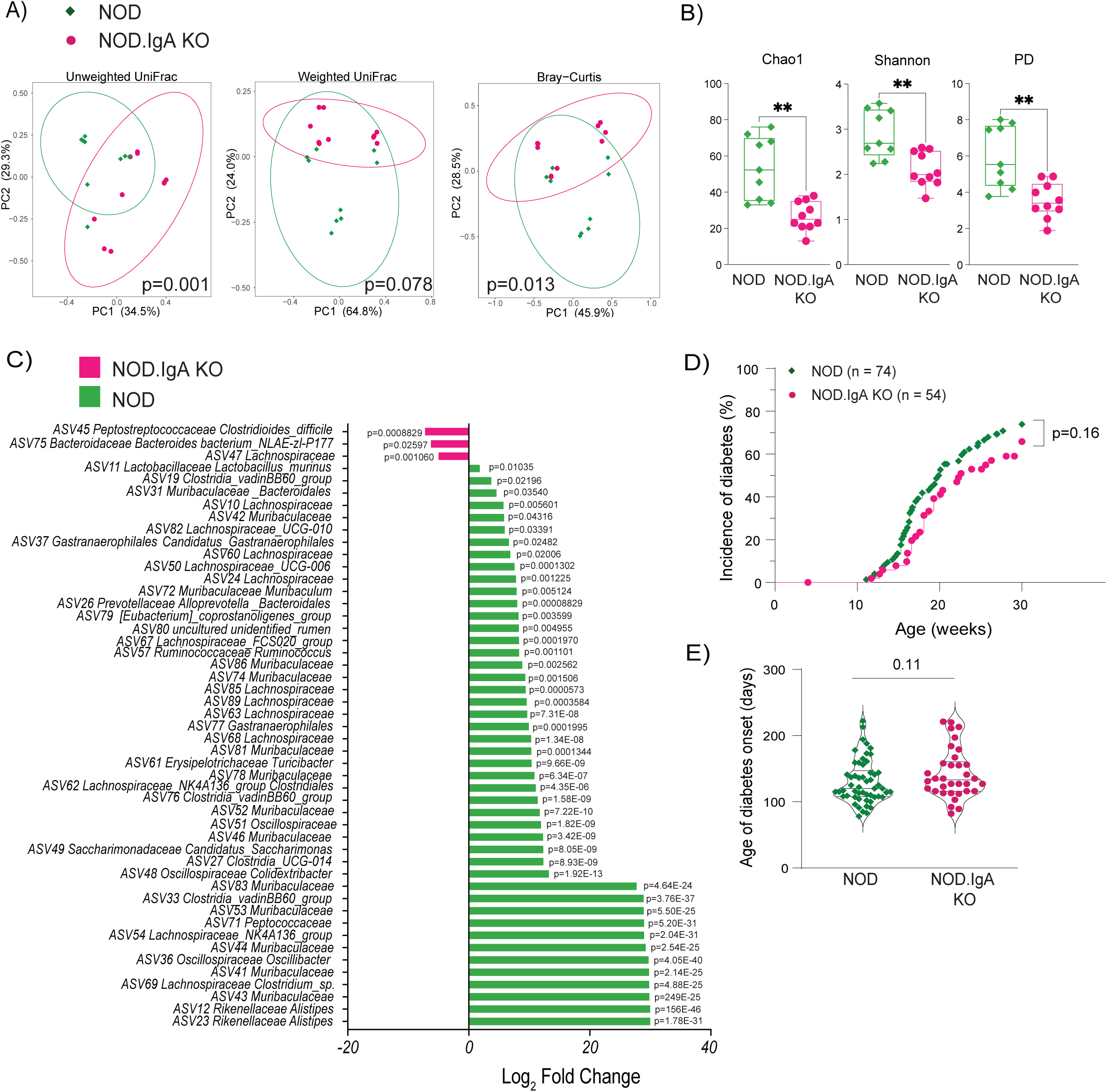
Multi-generational IgA deficiency reduces microbiome diversity without altering disease incidence. (A) Principal component analysis (PCA) of NOD-(green diamonds) and NOD.IgA KO-(pink circles) isolated fecal microbiota from mice at 12 weeks of age. (B) Box and whisker plots showing the alpha diversity of the fecal microbiota from both female cohorts. (C) Bar graph showing taxa enriched in the fecal material from NOD (green) and NOD.IgA KO (pink) cohorts. (D) T1D incidence in NOD (green diamonds) and NOD.IgA KO (pink circles) females over 30 weeks. (E) Age at disease onset in each of the two cohorts. *Data in (B) indicates mean, with error bars indicating min and max values. Violin plots in (E) indicate median and quartiles. For panels (A),(B),(D) & (E), each data point represents one mouse. p values were calculated using two-tailed Mann-Whitney U test for panels (A)-(C) and (E); Log-rank test was used for panel (D). **p<0.01*.

Since the NOD model is sensitive to microbiome changes^37^, we questioned whether IgA deficiency-induced dysbiosis would alter T1D incidence. We followed female NOD and NOD.IgA KO mice for glycosuria until 30 weeks of age. In spite of marked microbiome differences, disease incidence in NOD females (73.0%, 54/74) was similar to NOD.IgA KO females (63.0%, 34/54) (FIGURE 1D). The age at disease onset was also similar between the two cohorts (FIGURE 1E), suggesting that IgA deficiency neither exacerbated nor suppressed T1D development.

### Maternal IgA deficiency confers dominant resistance to T1D in IgA-sufficient offspring

Since IgA is a major humoral immune factor in breast milk, we next asked whether maternal provision of IgA had any bearing on offspring diabetes development. We designed a breeding scheme whereby NOD and NOD.IgA KO dams birthed IgA-sufficient offspring of the same genotype (FIGURE 2A). This scenario, where two genetically-identical offspring cohorts were birthed by genetically- and phenotypically-distinct dams, provided an opportunity to explore the influence of the maternal factors on offspring microbiome establishment, immune development and subsequent T1D pathogenesis. Offspring arising from NOD dams were termed “maternal IgA+ NOD.IgA HET” (mIgA+ HET), and those from NOD.IgA KO dams were termed “maternal IgA-NOD.IgA HET” (mIgA-HET).

**FIGURE 2.**
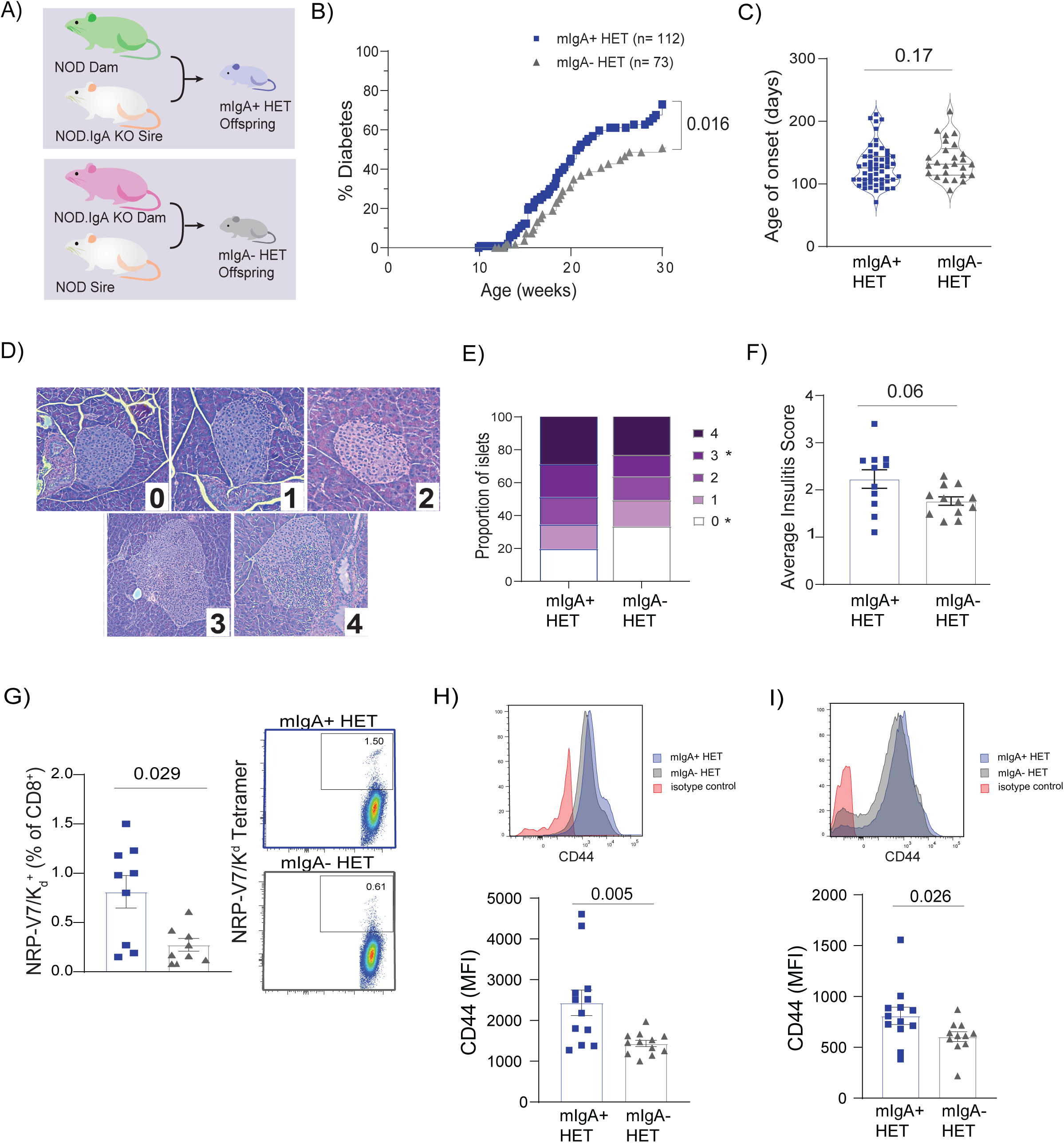
Maternal IgA deficiency confers dominant resistance to T1D in IgA-sufficient offspring. (A) Breeding scheme; NOD.IgA HET offspring from IgA-sufficient dams, referred to as mIgA+ HET (blue) were generated from NOD females crossed with NOD.IgA KO males. NOD.IgA HET offspring from IgA-deficient dams, referred to as mIgA-HET (grey) were generated from NOD.IgA KO females crossed with NOD males. (B) T1D incidence curve of mIgA+ (blue squares) vs mIgA-(grey triangles) NOD.IgA HET offspring. (C) Violin plot shows age of onset of T1D in each of the HET cohorts. (D) Insulitis scoring scale, with representative images of each score. (E) Insulitis scores of 12-13 week old pre-diabetic mIgA+ (blue squares) and mIgA- (grey triangles) HET mice, displayed as average score per mouse. (F) Insulitis scores displayed as average proportion of islets receiving each score. (G) Assessment of NPR-V7-reactive (IGRP mimitope) CD8^+^ T cells in the PLN at 12 weeks of age, expressed as percentage of total CD8^+^ population, with representative FACS plots. (H) Mean fluorescence intensity (MFI) of CD44 expression in PLN CD4^+^ populations at 12 weeks of age, with representative histograms. (I) MFI of CD44 expression in PLN CD8^+^ populations at 12 weeks of age, with representative histograms. *p-values were determined using the Log-rank test for panel (B), and the Mann-Whitney U test for all other plots. Each data point represents one mouse. Data in panels (G)-(I) are representative of 3 independent. In all panels, blue squares represent mIgA+ HET mice; grey triangles represent mIgA-HET mice.* p<0.05*.

mIgA- HET offspring had a decreased incidence of T1D, although age at onset was unaltered (FIGURE 2B,C). Interestingly, this reduced T1D incidence was not observed in IgA-deficient offspring from NOD.IgA KO dams (supplemental FIGURE 1), suggesting that maternal factors from an IgA-deficient dam reduce T1D susceptibility only when offspring have the ability to produce endogenous IgA. Consistent with lower T1D incidence, we observed reduced insulitis in adult pre-diabetic mIgA- HET females compared to age-matched mIgA+ HET females (FIGURE 2D-F). Furthermore, the frequencies of autoreacitve IGRP_206-214_ -specific CD8^+^ T cells were reduced in the pancreatic draining lymph nodes (PLN) of mIgA- HET mice (FIGURE 2G), along with lower activation in PLN T cell populations overall (FIGURE 2H,I). Collectively, these data suggest that maternal IgA deficiency suppresses autoreactive T cells and T1D development in genetically prone mice.

### Maternal IgA deficiency and associated microbiome changes induce a transient microbiota shift in IgA-sufficient offspring

The maternal microbiome has a strong influence on neonatal microbiome establishment and development. Both birth mode (vaginal birth vs cesarean section) and infant diet (breast milk vs formula) shape the colonizing microbiota during infancy^26,62–64^ in part through provision of microorganisms originating from breast milk as well as the maternal vaginal and GI tracts^62,63,65^. To test whether maternal IgA deficiency-linked protection is mediated by changes in the neonatal microbiome, we next determined whether the microbiome differences noted between IgA-sufficient and -deficient dams were propagated to their respective IgA-sufficient offspring. Furthermore, we examined how offspring microbiome composition evolved as the pups matured from the early life period (3 weeks of age) to pre-diabetic adulthood (12 weeks of age) in an effort to understand whether early differences were retained into adulthood.

Through these comparisons, we observed that the early (3 weeks) microbiota of each NOD.IgA HET cohort showed similarities to the maternal microbiota that contributed to differences in bacterial composition (FIGURE 3A). Alpha diversity in both cohorts was limited (FIGURE 3B), likely due to the immaturity of the developing microbiome at this life stage. Despite this, a number of taxa showed enrichment in one offspring cohort over the other (FIGURE 3C,D). Maternal transfer of microbes was evidenced in the enrichment of several species in the mIgA+ HET cohort, many belonging to the *Muribaculaceae* family. Bacterial families *Eubacteriaceae*, *Ruminococcaceae* and *Lactobacillaceae* were enriched in the mIgA- HET cohort, with a concomitant reduction of *Tannerellaceae* abundance (FIGURE 3C). At the species level, mIgA- HET pups showed significant enrichment of *Lactobacillus murinus*, while *C. difficile* and *Akkermansia muciniphila* were significantly enriched in mIgA+ HET pups (FIGURE 3D). Notably, none of the three species present at higher abundance in the NOD.IgA KO dams were enriched in the mIgA- HET offspring, possibly due to reduced cross-feeding activity in an altered ecosystem or these species being suppressed by the pups’ own antibody responses.

**FIGURE 3.**
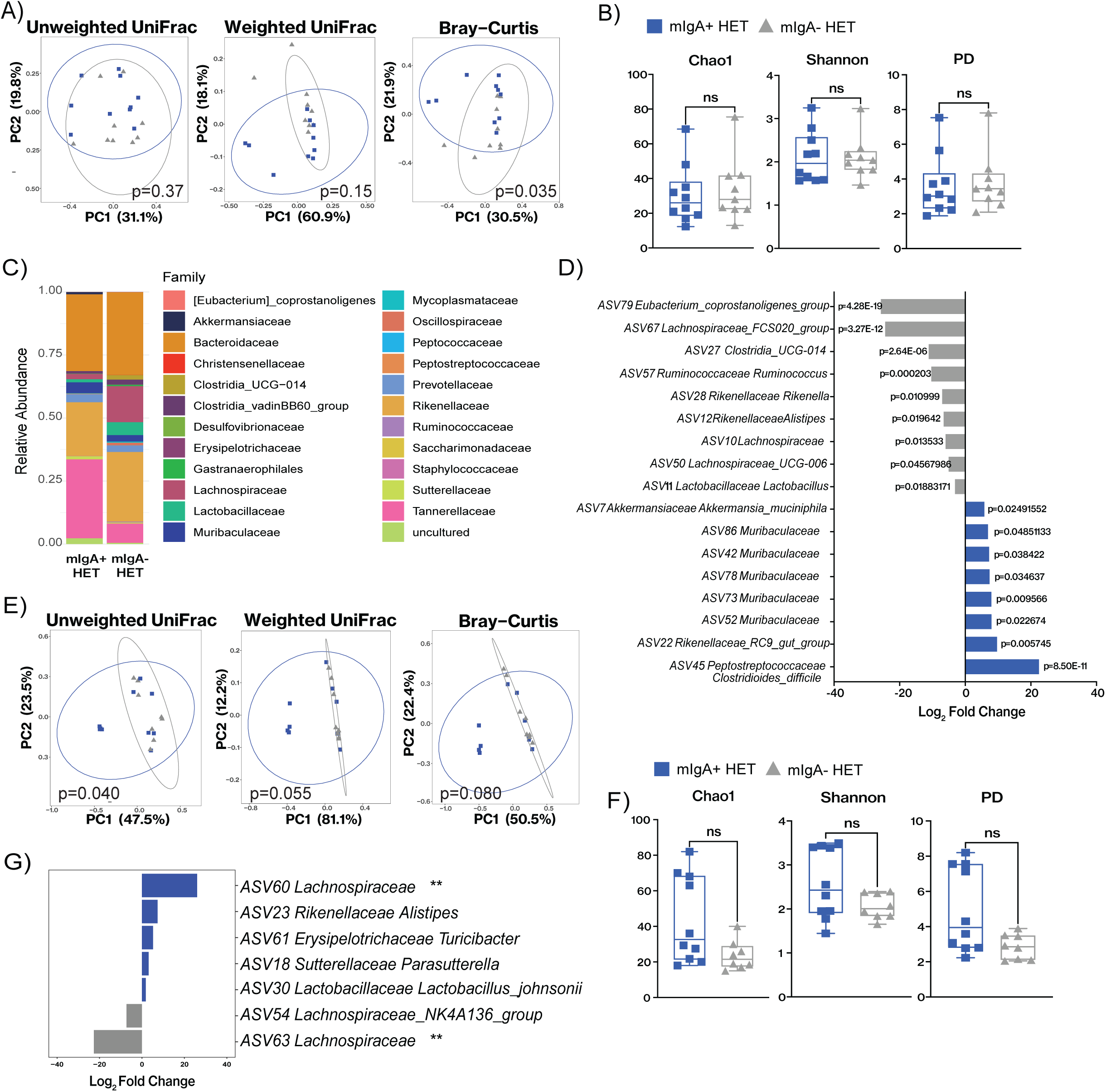
Maternal IgA deficiency and associated microbiome changes induce a transient microbiota shift in IgA-sufficient offspring. (A) PCA of mIgA+ HET (blue squares) and mIgA- HET (grey triangles) fecal microbiota at 3 weeks of age. (B) Alpha diversity of the two NOD.IgA HET cohorts at 3 weeks of age using various diversity measures. (C) Stacked bar graphs showing microbiota composition at the family taxonomic level in 3 week old HET cohorts. (D) Bar graph showing enriched taxa in each NOD.IgA HET cohort at 3 weeks. (e-h) Analysis of adult pre-diabetic (12 weeks) mIgA+ HET (blue squares) and mIgA- HET (grey triangles) fecal microbiota, including (E) PCA, (F) alpha diversity plots, and (G) bar graph showing taxa enrichment. *Data in (B)and (F) indicates mean, with error bars indicating min and max values. p-values were determined using the unpaired t test. Each data point represents one mouse. ns: non-significant; **p<0.01*.

The compositional differences between the offspring cohorts at 3 weeks were largely lost in adulthood (12 weeks). While four littermates of the mIgA+ HET cohort showed increased heterogeneity that could stem from litter effects, the microbiome composition in the majority of mIgA+ HET mice were similar to mIgA- HET offspring (FIGURE 3E,F). When the four deviant littermates were removed from analysis, only two species showed significantly altered abundance between the two cohorts (FIGURE 3G). This finding is in line with other studies on the early microbiome, where sub-optimal GI events during the weaning period led to dysregulated immunity in adulthood without leaving a detectable mark on the adult microbiota^21,22^.

Altogether, our findings suggest that maternal IgA deficiency induced a transient microbiome alteration that was sufficient to mediate long term resilience to T1D despite the absence of persisting dysbiosis in adulthood.

### IgA-sufficient offspring from dysbiotic IgA-deficient dams harbor a significantly altered metabolome

Host and microbes interact and shape the metabolome in the gut, and the presence or absence of key genera can strongly influence the metabolome composition^66^. This is true for mIgA+ and mIgA- HET pups, which displayed marked differences in their gut metabolome at 5 weeks of age, when composition was no longer influenced by breast milk (supplemental FIGURE 2).

PCA revealed that the metabolome of each cohort clustered based on maternal origin (FIGURE 4A), displaying a number of metabolites significantly altered between the two cohorts (FIGURE 4B). We first compared the abundance of metabolites with known immunomodulatory properties. Short chain fatty acids (SCFA) including acetate, propionate and butyrate were all decreased in the diabetes-resistant mIgA- HET cohort (FIGURE 4C). Similarly, all identified indole derivatives (products of tryptophan metabolism and agonists of the aryl hydrocarbon receptor, [AhR]) were decreased in mIgA- HET mice (data not shown). While SCFA and indoles are typically associated with positive health outcomes, the lower levels observed are consistent with a more dysbiotic microbiome. Several pro-inflammatory lipid mediators, including prostaglandins B2, C2, H2, I2 and leukotrienes B4 & E4, were detected at lower concentrations in the mIgA- HET cohort (FIGURE 4D and data not shown). Of the metabolites that were increased in the mIgA- HET samples, many were molecules involved in amino acid metabolism. Indeed, pathway analysis using MetaboAnalyst 5.0^67^ identified significant alterations in amino acid metabolism and folate metabolism (FIGURE 4E). The identified pathways were further dissected with MetOrigin^68^ to evaluate host, microbiota and co-metabolic influences, which corroborated multiple pathways associated with the metabolism of amino acids in these three compartments, as well as pathways related to inflammation (FIGURE 4F). Importantly, in both the microbiota and co-metabolism compartments, but not in the host, pathways related with SCFA and folate metabolism were significantly altered, with various pathway intermediates showing differential abundance. Small differences in retinol acid metabolism, important for the induction of IgA production, were classified as solely associated with altered host metabolism, and microbial metabolic pathways were responsible for differences in aromatic hydrocarbon degradation.

**FIGURE 4.**
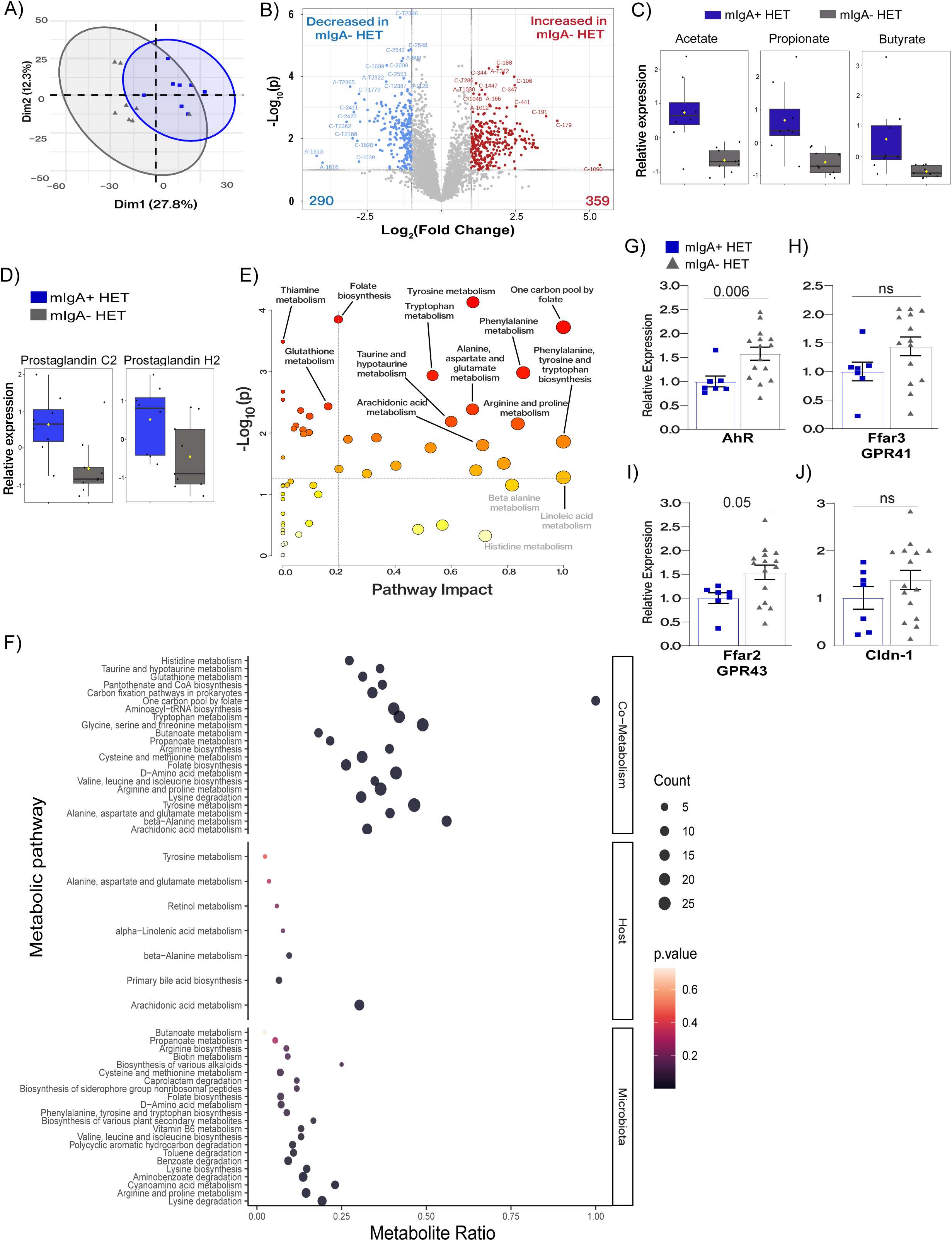
IgA-sufficient offspring from dysbiotic IgA-deficient dams harbor a significantly altered metabolome. (A) PCA of fecal metabolome of 5 week old mIgA+ (blue square) and mIgA- (grey triangle) HET mice. (B) Volcano plot showing metabolites that are up-regulated (red; 359 metabolites) and down-regulated (blue; 290 metabolites) in the mIgA- HET cohort vs mIgA+ HET pups. (C) Comparison of fecal short chain fatty acids and (D) select prostaglandin levels between mIgA+ and mIgA- HET mice. (E) Pathway analysis showing the impact of metabolite abundance differences on metabolic pathways using MetaboAnalyst 5.0. (F) Pathway analysis showing altered metabolic pathways separated into microbiome, host and co-metabolism compartments. (g-j) Relative comparison of (G) *AhR*, (H) *Ffar3* (GPR41), (I) *Ffar2* (GPR43) and (J) *Cldn-1* mRNA levels in 4-5 week old mIgA+ (blue square) and mIgA- (grey triangle) HET mice. *Data is shown as means ± SEM. p-values were determined using the Mann-Whitney U test. Each data point (panels a, c, d, g-j) represents one mouse*.

Notably, host expression of AhR and SCFA receptors (GPR43 and GPR41) was elevated in the ileum of 5 week old mIgA- HET mice (FIGURE 4G-I), along with up-regulation of the gene encoding tight junction protein Claudin 1, *Cldn1*^69^, in some members of the mIgA- HET cohort (FIGURE 4J). Thus, maternal IgA deficiency contributed to significant changes in the early metabolome of the offspring and to amplified expression of receptors for immunomodulatory metabolites, enhancing the ability of the offspring to sense the microbiome. Many of the metabolomic differences were indicative of an increased inflammatory environment along the GI tract in the mIgA- HET cohort in the early post-weaning period.

### IgA sufficient offspring from dysbiotic mothers show a more transcriptionally active GI environment at weaning

The striking difference in disease incidence between the NOD.IgA HET offspring from IgA-sufficient vs -deficient dams suggests there are maternal factors at work during the neonatal period that either promote or dampen efficient immune regulation. Immune tolerance is established largely during a pre-weaning “window of opportunity”, where host interaction with the microbiota results in the differentiation and proliferation of regulatory CD4^+^ T lymphocyte populations^21,22^. Hallmarks of an appropriately robust response to the microbiota during this period include the proliferation of immune cell populations, expansion of the adaptive immune repertoire, and the induction of immune-related gene expression along the GI tract^22,70–72^. In short, this ‘weaning reaction’^22^ involves the induction of GI immune responses that would generally be considered pro-inflammatory but are in fact essential for health.

Since pups are weaned at 21 days of age and others have reported the weaning reaction occurring within the days following weaning^22^, we performed RNAseq on ileal (small bowel, SB) and colonic tissue (large bowel, LB) from 25 day old pups. In the LB tissue, only 9 transcripts were differentially expressed between the two cohorts (Supplemental FIGURE 3). We focused our analysis on the SB tissue, where we found 47 ileal transcripts exhibiting altered expression, all of them showing up-regulation in the mIgA- HET pups (FIGURE 5A,B). Most notably, with the possible exception of one transcript (*Upp1*), all up-regulated genes have known or predicted roles in host immune response. Gene ontology (GO) analysis identified host antimicrobial immune responses as dominant pathways up-regulated in mIgA- HET SB (FIGURE 5C). Specifically, ileal tissues from mIgA- HET pups exhibited a gene signature of up-regulated pattern recognition receptor (PRR) activation & signaling, cytokine & chemokine expression, antigen processing & presentation, and interferon (IFN) responses, which are characteristic of a robust weaning reaction in response to colonization^22,28^ (FIGURE 5D). Similarly, quantitative RT-PCR on ileal tissues from a larger number of samples (n=9 mIgA+ HET; n=15 mIgA- HET) at 25-30 days of age showed elevated *il10*, *ifng*, and *lcn2* (FIGURE 5E-G). These data collectively suggest that an amplified weaning reaction underscores the T1D resistance seen in the mIgA- HET cohort.

**FIGURE 5.**
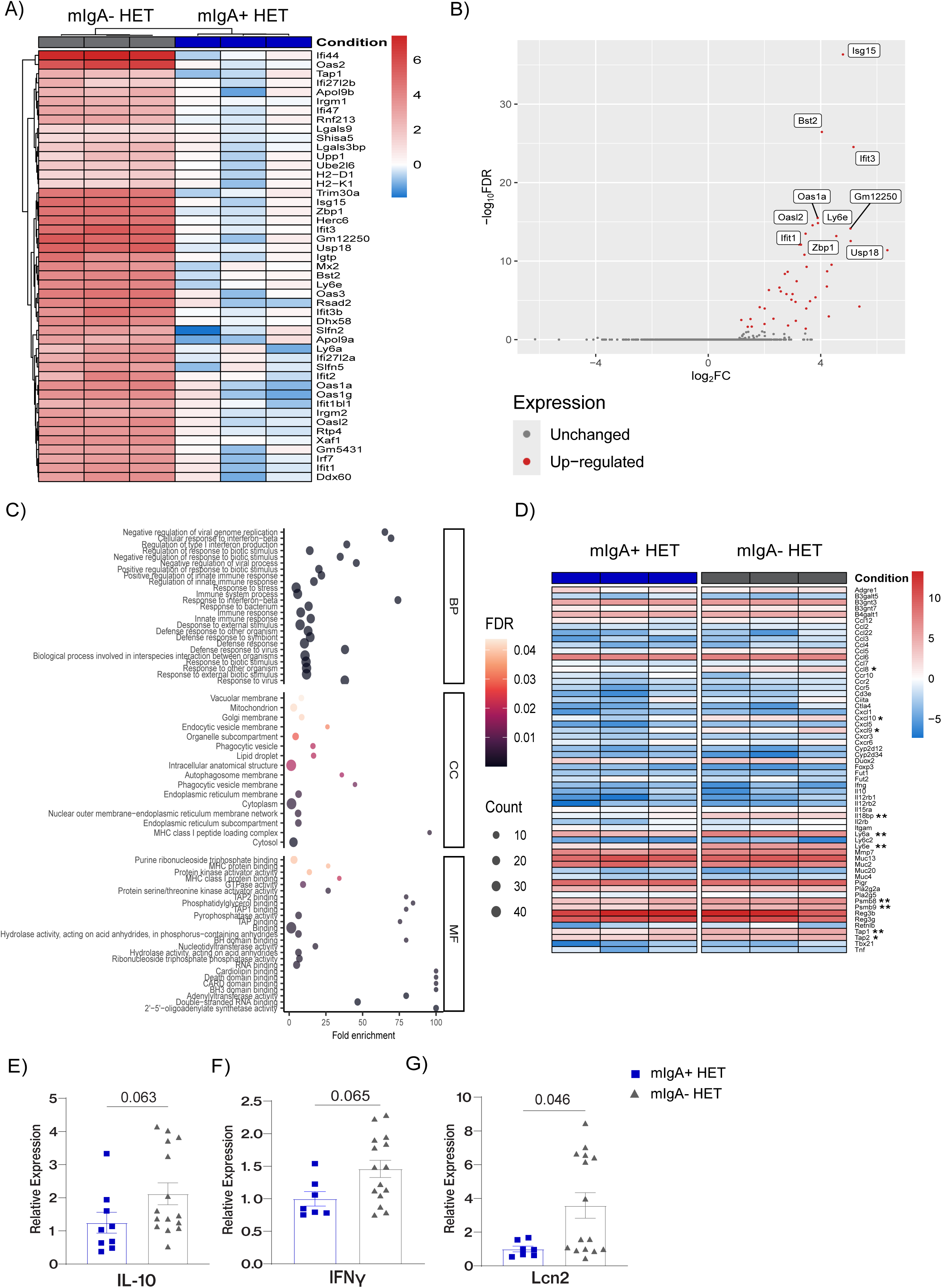
IgA sufficient offspring from dysbiotic mothers show a more transcriptionally active GI environment at weaning. (A) Heatmap and (B) volcano plot displaying ileal transcripts (FDR<0.05) with altered expression in the mIgA- HET cohort at 25 days of age, as compared to the mIgA+ HET cohort (n=3 each). (C) Gene Ontology (GO) analysis of pathways and processes affected by up-regulated transcripts in the ileum, where the top 20 GO terms are displayed. (D) Heatmap showing relative expression of weaning reaction-associated genes. (e-g) Quantitative RT-PCR results showing relative expression of indicated transcripts in ileal tissue from 25-30 day old mIgA+ HET (blue square) and mIgA- HET (grey triangle) pups. *BP: biological process; CC: cellular compartment; MF: molecular function. IL-10: interleukin-10; IFNγ: interferon gamma; Lcn2: lipocalin 2; TGFβ: tumor growth factor beta; IL-6: interleukin-6. Data in panels (E)-(G) shown as means ± SEM. p-values were determined using the Mann-Whitney U test. Each data point represents one mouse. * p<0.05; ** p<0.01*.

### Maternal dysbiosis promotes heightened mucosal inflammation and IgA immunity in offspring during early life

To assess the effect of the weaning reaction, we profiled the immune cell populations in the LB, SB and mesenteric lymph nodes (MLN) at 3 and 4 weeks of age in each HET cohort. Maternal IgA deficiency marginally altered the B cell compartment in the SB and LB (FIGURE 6A), but promoted increased IgA class-switching (FIGURE 6B) and costimulatory molecule CD86 expression (FIGURE 6C). Within the plasma cell (PC) population, the mIgA- HET cohort showed a higher proportion of IgA-producing PC with a greater overall IgA staining intensity, indicating increased IgA production per cell (FIGURE 6D). The increased IgA class switching was accompanied by higher frequencies of Tfh cells in the MLN (FIGURE 6E) and an amplified germinal center reaction at 3 weeks of age (FIGURE 6F), suggesting that maternal IgA deficiency promoted T-dependent IgA responses.

**FIGURE 6.**
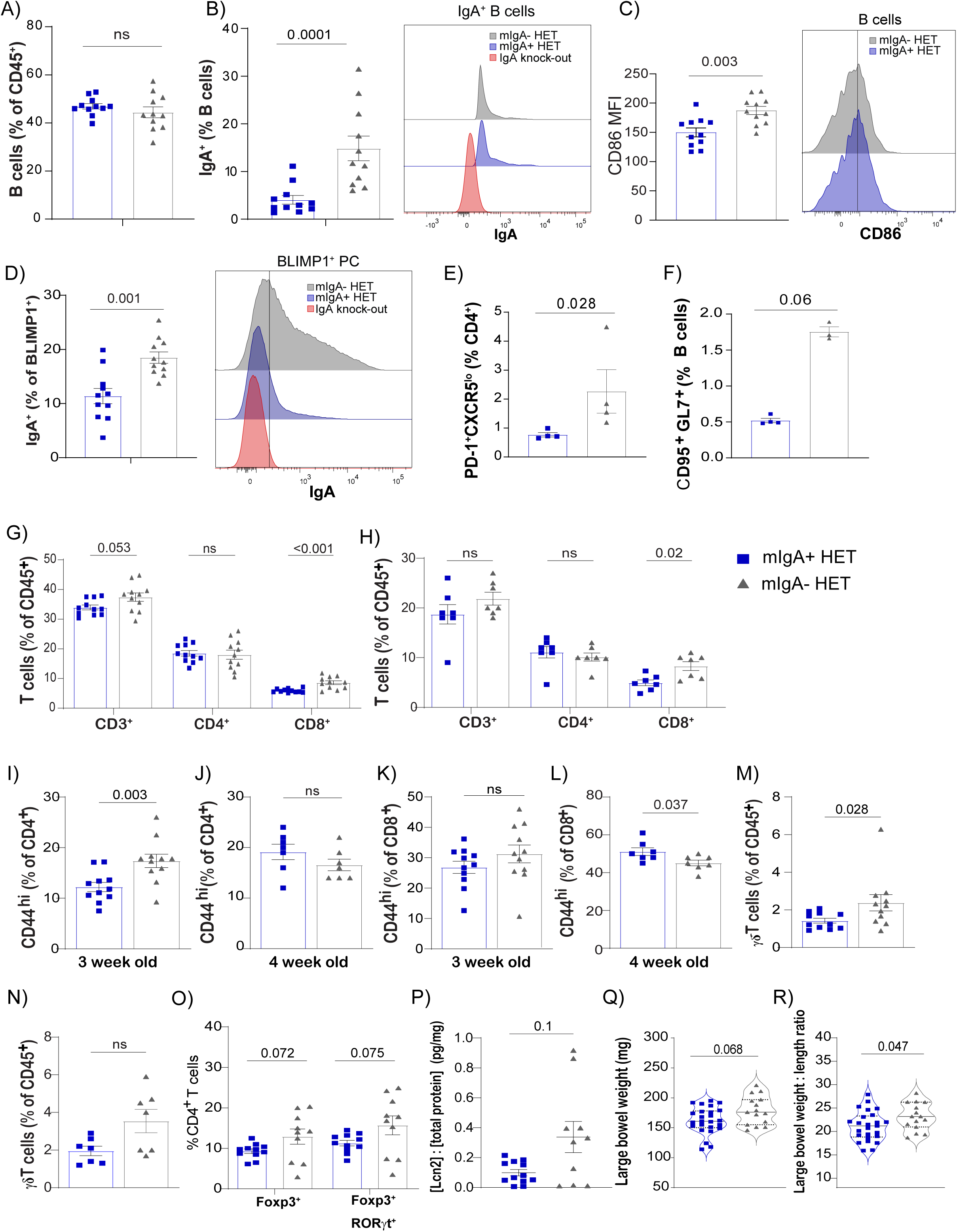
Maternal dysbiosis promotes heightened mucosal inflammation and IgA immunity in offspring during early life. Flow cytometry analysis of GI immune cell populations at 3 weeks of age, with bar graphs and representative FACS plots, where mIgA+ HET data is indicated in blue squares and mIgA- HET data is indicated in grey triangles. (A) Total B cells and (B) IgA^+^ B cells in the LB, with representative histograms. (C) Mean fluorescence intensity (MFI) of B cell CD86 expression in the LB, with representative histograms. (D) IgA^+^ plasma cells in LB, with representative histograms. (E) Follicular helper T cells and (F) germinal center B cells in MLN of 3 week old HET mice. Total T cells, CD4^+^, & CD8^+^ T cell populations in the (G) LB and (H) SB of HET mice at 4 weeks of age. CD44 expression on SB CD4^+^ T cells at (I) 3 weeks and (J) 4 weeks of age. CD44 expression on SB CD8^+^ T cells at (K) 3 weeks and (L) 4 weeks of age. γδ T cells in SB at (M) 3 weeks and (N) 4 weeks of age. (O) FoxP3^+^ and FoxP3^+^RORγt^+^ regulatory CD4^+^ T cells in the LB of HET mice at 3 weeks of age. (P) Fecal Lcn-2 protein levels in 4-5 week mice, expressed as ratio of total fecal protein. (Q) Colonic weight and (R) weight:length ratio in 4-5 week old mice. *For all panels, data from mIgA+ HET mice is indicated in blue squares, mIgA- HET mice in grey triangles. Data is shown as means ± SEM for all data. p-values were determined using the Mann-Whitney U test. Each data point represents one animal. Data in all panels are representative of ≥ 2 independent experiments*.

The heightened immune phenotype was not restricted to the B cell compartment. While the T lymphocyte compartments in the LB and SB of mIgA+ and mIgA- HET cohorts were largely similar (FIGURE 6G,H), we noted an increase in activated CD4^+^ T cells in the SB at weaning which normalized by 4 weeks of age (FIGURE 6I,J). Similar trends of suppressed activation post-weaning were observed for SB CD8^+^ T cells (FIGURE 6K,L) and γδ T cells (FIGURE 6M,N), supporting a transient overall activated state at weaning that dissipates henceforth. One previously reported key outcome of the weaning reaction is the differentiation of inducible regulatory T cells (iTregs, CD4^+^FOXP3^+^ROR𝛾t^+^)^22^, which critically mediate immune tolerance to both dietary antigens and luminal microbial presence. We assessed SB and LB of mIgA+ and mIgA- HET pups for frequencies of iTregs, and observed that maternal IgA deficiency increased the frequency of ROR𝛾t^+^ iTregs in the LB (FIGURE 6O). Thus, we report that maternal IgA deficiency fosters changes in the neonatal GI environment that potentiate a strong weaning reaction and the differentiation of iTregs to curb T1D development.

Changes in LB immune populations prompted us to further evaluate the colon. Indications of colonic inflammation include increased fecal Lipocalin-2 (Lcn-2) abundance^73^ and elevated colonic weight:length ratio. When compared to their mIgA+ HET counterparts, newly weaned mIgA- HET pups exhibited these symptoms of increased GI inflammation (FIGURE 6P-R). Interestingly, the elevated colonic weight:length ratio was affected not by colonic shortening, as is the case in colitis, but by increased weight of the colon, indicating that the immune processes at work during early life are different from those that contribute to inflammatory diseases in adulthood.

### Maternal IgA deficiency regulates gut barrier function and GI humoral responses later in life

Both human^40^ and animal studies^41^ have shown evidence of a defective mucus layer in T1D. To test whether T1D resistance in mIgA- HET pups was associated with enhanced gut barrier function later in life, we carried out histological examination of the colonic mucus layer in the two NOD.IgA HET cohorts. At 12 weeks of age, we observed a strengthened colonic mucus layer in the mIgA- HET cohort, with enhanced integrity and thickness (FIGURE 7A). Although there was no evidence of bacterial encroachment directly on the epithelial barrier in either cohort, the reduced thickness of the mucus layer in the mIgA+ HET LB resulted in closer proximity of the microbes to the barrier (FIGURE 7B).

**FIGURE 7.**
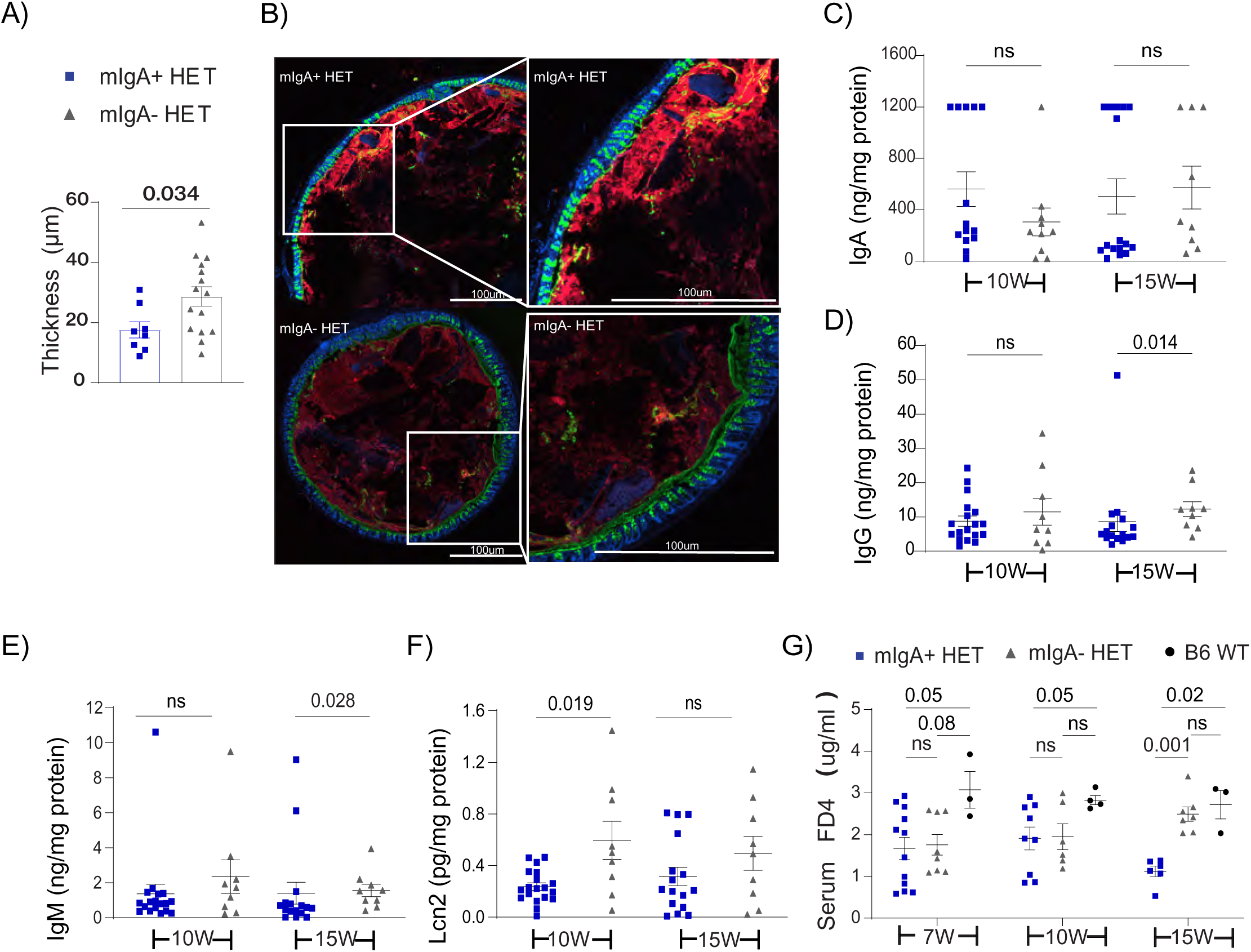
Maternal IgA deficiency regulates gut barrier function and GI humoral responses later in life. (A) Mucus layer thickness in colon specimens of pre-diabetic mIgA+ and mIgA- HET mice at 12-13 weeks of age. (B) Representative cross-sectional images of the colon showing bacteria (red), mucus (lectin, green) and intestinal epithelial layer (dapi, blue). Fecal levels of (C) IgA, (D) IgG, (E) IgM and (F) Lcn-2 in pre-diabetic 12-14 week mIgA+ and mIgA- HET mice. (G) Gut permeability in mIgA+ and mIgA- HET mice at 7, 10 & 15 weeks of age, compared to age-matched C57BL/6 mice (black circles). *For all panels, data from mIgA+ HET mice is indicated in blue squares, mIgA- HET mice in grey triangles. Data is shown as means ± SEM. p-values were determined using the Mann-Whitney U test. Each data point represents one mouse*.

We next measured fecal Ig and Lcn-2 content in mIgA+ and mIgA- HET mice at 10 and 15 weeks of age as a readout for altered immune responses to a potentially breached GI barrier. Fecal total IgA was highly variable in both cohorts (FIGURE 7C), while fecal IgG, IgM, and Lcn-2 showed increased abundance in mIgA- HET mice (FIGURE 7D-F). Furthermore, gut permeability assays with orally gavaged FITC-dextran (FD4) beads did not find altered paracellular barrier function in NOD.IgA HET mice (FIGURE 7G), suggesting that the increased intestinal IgG production in the adult mIgA- HET cohort was not caused by heightened immune activation due to a compromised gut barrier.

### Heightened disease susceptibility in offspring of IgA-sufficient dams is attenuated by fostering to IgA-deficient dams

To disentangle the contributions of pre- and peri-natal maternal influences from those that are enacted postnatally, we conducted fostering experiments within 24 hours of birth. NOD.IgA HET pups born to NOD dams were fostered to newly post-partum NOD.IgA KO dams, and the resultant pups monitored for diabetes onset until 30 weeks of age. NOD.IgA HET pups born to NOD dam but fostered to NOD.IgA KO dam (henceforth referred to as F^KO^ mIgA+ HET) displayed reduced T1D incidence compared to NOD.IgA HET pups born to NOD dam (mIgA+ HET) who were not fostered, while mIgA+ HET pups fostered to NOD dams (F^WT^ mIgA+ HET) did not exhibit blunted T1D (FIGURE 8A). Notably, the level of protection in fostered progeny did not reach that exhibited by mIgA- HET pups, suggesting important maternal influences during the pre- and peri-natal periods. Thus, postnatal factors are a significant, though not exclusive, contributor to maternal IgA deficiency-linked disease resistance.

**FIGURE 8.**
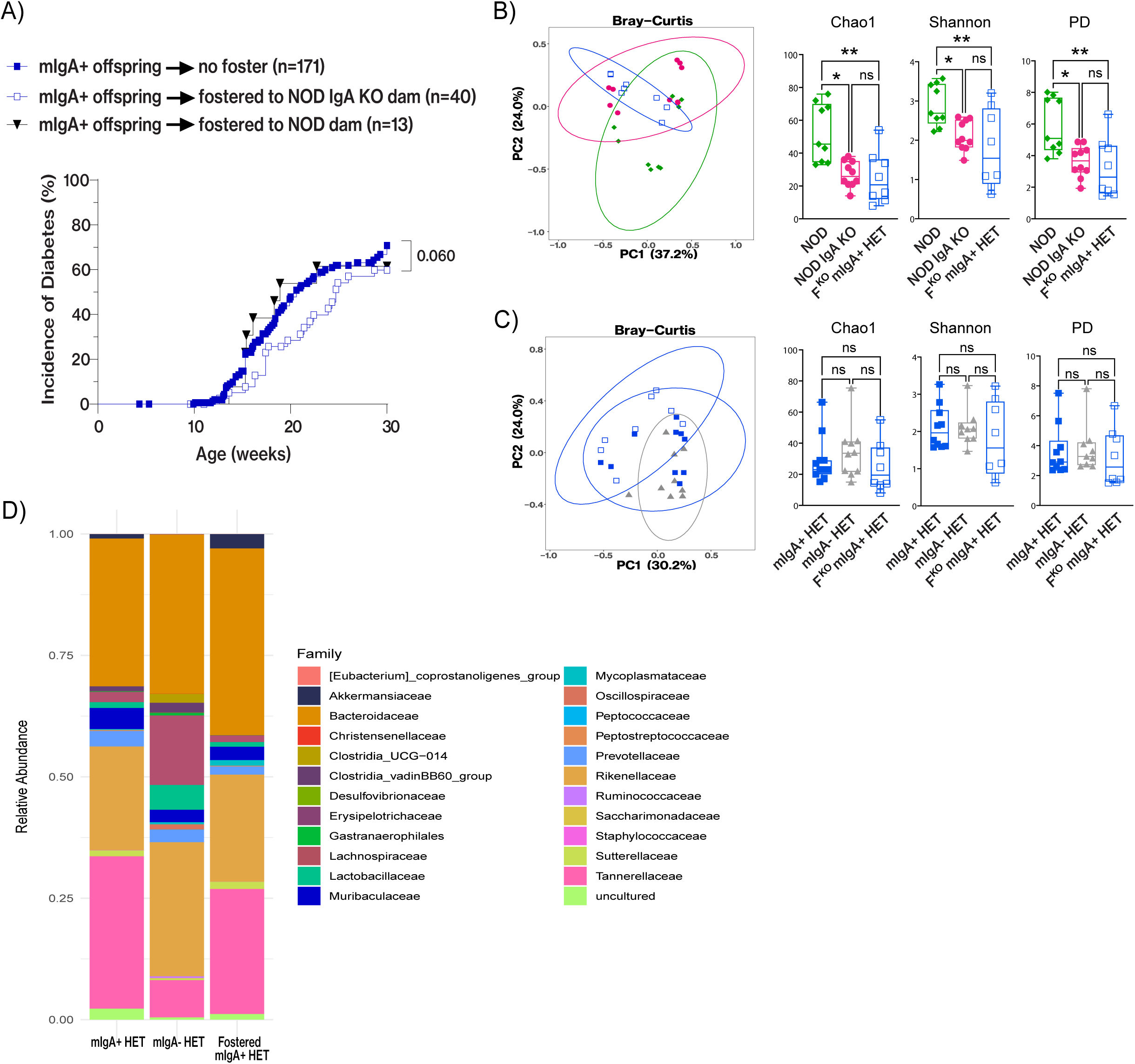
Heightened disease susceptibility in offspring of IgA-sufficient dams is attenuated by fostering to IgA-deficient dams. (A) Disease incidence curve for fostered offspring, showing T1D incidence of NOD dam-born IgA-sufficient offspring fostered to NOD.IgA KO dams (open blue squares), in comparison to those not fostered (solid blue squares) or fostered to another NOD dam (black triangle). Microbiome analysis of fostered mIgA+ HET pups at 3 weeks of age, with comparisons to (B) the birth dam (NOD) and foster dam (NOD.IgA KO) cohorts, and (C) the birth sibling (mIgA+ HET) and foster sibling (mIgA- HET) cohorts. (D) Stacked bar graph showing microbiota composition at the family taxonomic level for fostered mIgA+ HET pups (left), mIgA+ HET pups (middle) and mIgA- HET pups (right). *For all panels, fostered mIgA+ HET data is indicated in open blue squares, mIgA+ HET data is indicated in solid blue squares, mIgA- HET data is indicated in grey triangles. For panel (A) p-value was determined using Gehan-Breslow-Wilcoxon test. Data in (B) and (C) is shown as means with error bars indicating min. and max. values. p-values were determined using Mann-Whitney U test. Each data point represents one mouse*.

F^KO^ mIgA+ HET mice had early exposure to microbes from two maternal sources harboring diverse microbial communities, prompting the question of whether and how birth vs fostering dams influenced the pup microbiome. 16S rRNA gene sequencing of fecal samples collected from fostered progeny at 3 weeks of age showed that the fostered pups more closely resembled the foster dam (NOD.IgA KO) than the birth dam (NOD) with respect to both alpha and beta diversity (Bray-Curtis) (FIGURE 8B). Despite this, the early microbiome in these pups appeared more similar to mIgA+ HET pups (ie; their biological littermates) than mIgA- HET pups (ie; their “adopted” littermates) (FIGURE 8C).

Furthermore, rather than being a composite makeup of the two HET cohorts, the fostered offspring harbored a unique microbiota composition reflective of their unique early life period. Of note, F^KO^ mIgA+ HET pups harbored significantly increased abundance of both *C. difficile* and *A. muciniphila* compared to the mIgA- HET pups. This phenotype remarkably mirrored what was seen in diabetes-susceptible mIgA+ HET pups (FIGURE 3C), with both of these species associating positively with elevated T1D incidence. Additionally, F^KO^ mIgA+ HET pups harbored increased *Parabacteroides goldsteinii* abundance compared to mIgA- HET pups and more *Bacteroides sartorii* than mIgA+ HET pups. *Bacteroides acidifaciens* was depleted in fostered pups compared to both mIgA+ and mIgA- HET cohorts. Differences in composition were evident when evaluated at the family taxonomic level, although none reached significance (FIGURE 8D).

Taken together, the microbiome composition in fostered mIgA+ HET pups at 3 weeks of age set them apart from both their biological and foster siblings, reflecting their unique early life exposures. The most striking change noted between foster siblings was enrichment of *A. muciniphila* and *C. difficile* in the fostered mIgA+ HET pups, a pattern consistent with observations for mIgA+ vs mIgA- HET pups.

### IgA deficiency and resultant dysbiosis alter the breast milk proteome in lactating dams

IgA is the most abundant antibody in breast milk, followed by IgG and IgM^74^. Breast milk antibodies contribute to neonatal development in various ways, from providing passive immunity against infection^75,76^ to modulating the microbiome to inducing oral tolerance^77–79^. Comparing the Ig concentrations in breast milk samples collected from NOD and NOD.IgA KO dams, we noted that the most abundant Ig, IgA, was indeed absent in NOD.IgA KO dams (FIGURE 9A). The loss of IgA was accompanied by a compensatory rise in other antibody isotypes, showing an average 5.3-fold increase in IgG and 6.6-fold increase in IgM (FIGURE 9A).

**FIGURE 9.**
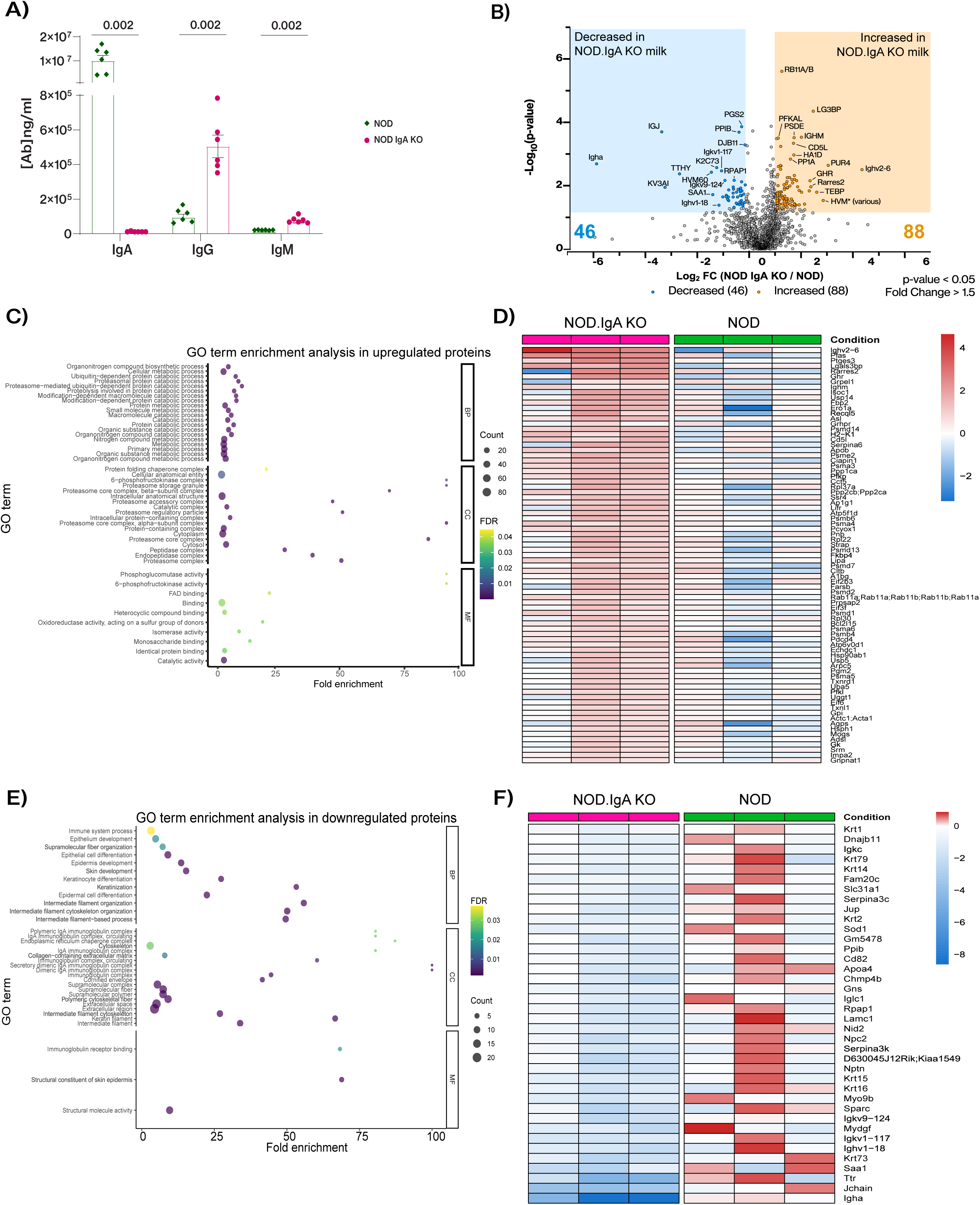
IgA deficiency and resultant dysbiosis significantly alter the breast milk proteome in lactating dams. (A) Immunoglobulin content of mouse breast milk collected from NOD (green diamonds) and NOD.IgA KO (pink circles) dams at post-partum day 10-12, assayed via ELISA. (B) Proteomic analysis of breast milk visualized using a volcano plot, indicating up- and down-regulated proteins in milk from NOD.IgA KO vs. NOD dams. (C) GO analysis and (D) heatmap of significantly up-regulated (FDR<0.01) proteins in milk collected from NOD.IgA KO vs. NOD dams. (E) GO analysis and (F) heatmap of down-regulated proteins (FDR<0.01) in milk collected from NOD.IgA KO vs. NOD dams. *Data in panel (A) is shown as means ± SEM. p- values were determined using Mann-Whitney U test. Each data point represents one mouse*.

We next performed proteomic analysis on breast milk samples collected from NOD and NOD.IgA KO milk (n=3 each, 10-12 days post-partum). We quantified 1982 proteins in total, with 1350 detected proteins shared between all samples. Using an unpaired t-test (p-value<0.05 and fold change > 1.5) we found 46 decreased and 88 increased proteins in abundance in the NOD.IgA KO milk (FIGURE 9B). Consistent with the ELISA data (FIGURE 9A), marked changes were observed in the antibodies of NOD.IgA KO milk. Ig heavy chain 𝛼 constant region and the J chain were depleted (Supplemental FIGURE 4A), and accompanied by an enrichment of the Ig heavy chain 𝜇 constant region (ie IgM heavy chain) in NOD.IgA KO milk (Supplemental FIGURE 4a). Ig γ heavy chain regions also showed marked alterations between the two cohorts, albeit with heterogeneous distribution (Supplemental FIGURE 4B). Significant reductions in the relative abundance of 𝝹 and 𝜆1 light chain constant regions were also seen (Supplemental FIGURE 4C). Several variable regions also showed differential abundance between the NOD and NOD.IgA KO milk samples, with IgH V2-6 showing the largest increase and IgH V1-18 and V3-6 showing a decrease in protein abundance (FIGURE 9B). Similarly, numerous 𝝹 light chain variable regions showed differential abundance between the two dam cohorts (FIGURE 9B). Thus, IgA deficiency imparts significant changes in antibody subclass and repertoire in the maternal milk.

Increased levels of other immunomodulatory proteins were identified in the NOD.IgA KO milk, including prostaglandin E synthase 3, chemerin (RARRES2), H-2K^d^ (MHC I), and CD5 antigen-like (CD5L). Additionally, myeloid-derived growth factor (MYDGF) and chemokine (C-X-C motif) ligand 17 (CXCL17) were found in decreased abundance in NOD.IgA KO milk. Pathway enrichment analysis with Enrichr^80^ identified several immune signaling pathways as highly increased in NOD.IgA KO milk, including Type I and III interferon responses, mTORC1 pathway, and Myc pathway targets. GO and KEGG pathway analyses indicated NOD.IgA KO milk was enriched in proteins involved in several cellular metabolic processes including biosynthesis of amino acids, biosynthesis of amino and nucleotide sugars, glycolysis, carbon metabolism, and pentose phosphate pathway (FIGURE 9B,C). Moreover, many proteasome subunit components (e.g. Psmd14, Psme2, Psma3, Psmb8, Psmb6, Psma4) were increased in abundance in NOD.IgA KO milk. In line with this, STRING analysis clearly showed substantial interactions between the various proteasome components (supplemental FIGURE 5A) and GO enrichment analysis indicated significant up-regulation of protein catabolic processes in these samples (FIGURE 9D). Breast milk is known to contain immune cells and secreted extracellular proteasome components^81^; however, since our extracted proteins encompass both intracellular and secreted products, it is unclear whether our data show increased secretion of extracellular proteasome or higher intracellular proteasome activity in milk from IgA-deficient dysbiotic dams.

Proteins significantly decreased in NOD.IgA KO milk included those involved in structural integrity and cytoskeleton organization (FIGURE 9E,F). STRING analysis identified associations between many of these proteins (Supplemental FIGURE 5B) and further research established their participation in extracellular matrix remodeling and differentiation of epithelial cells. Reduced levels of these proteins was predicted by Enrichr to strongly affect the process of epithelial mesenchymal transition, although if and how alterations in this pathway affect post-natal GI development remains to be better understood.

In summary, breast milk protein content differed substantially between the NOD and NOD.IgA KO dams. Dissimilarities in antibody subclass and variable region usage reflect significant changes in the antibody repertoire and effector function, prompting further investigation into the antigenic specificities of these antibodies. Increased cellular metabolic activity and immune signaling capacity are clear indicators of amplified potential for promoting a heightened immunogenic environment in the neonatal GI tract. Deeper investigation into the multi-factorial changes in breast milk, beyond protein composition, will shed additional light on the protective mechanisms at play.

### Milk antibodies from IgA-sufficient vs -deficient dams show differing potential to modulate offspring microbiota

Prompted by the dissimilar antibody repertoire in WT vs NOD.IgA KO dams, we next asked whether the milk from NOD vs NOD.IgA KO dams contained antibodies of dissimilar antigenic specificities, with differing abilities to modulate the neonatal microbiome. Fecal microbe populations from 3 week old mIgA+ HET and mIgA-HET pups was subjected to a binding assay with milk collected from 10-12 day postpartum NOD and NOD.IgA KO dams (FIGURE 10A). We found that nearly 60% of NOD milk IgA was capable of binding fecal microbes from mIgA+ HET pups, but showed less ability to coat microbes from mIgA-HET pups (FIGURE 10B). This is likely a result of the different microbial communities present within the two pup cohorts and mirrors the finding in humans that the maternal milk IgA repertoire is unique and tailored to a mother’s inherent offspring^82^. Furthermore, given the absence of IgA in the NOD.IgA KO dam milk, it is very likely that a substantial proportion of the mIgA-HET pup microbiota escaped modulation by milk antibodies.

**FIGURE 10.**
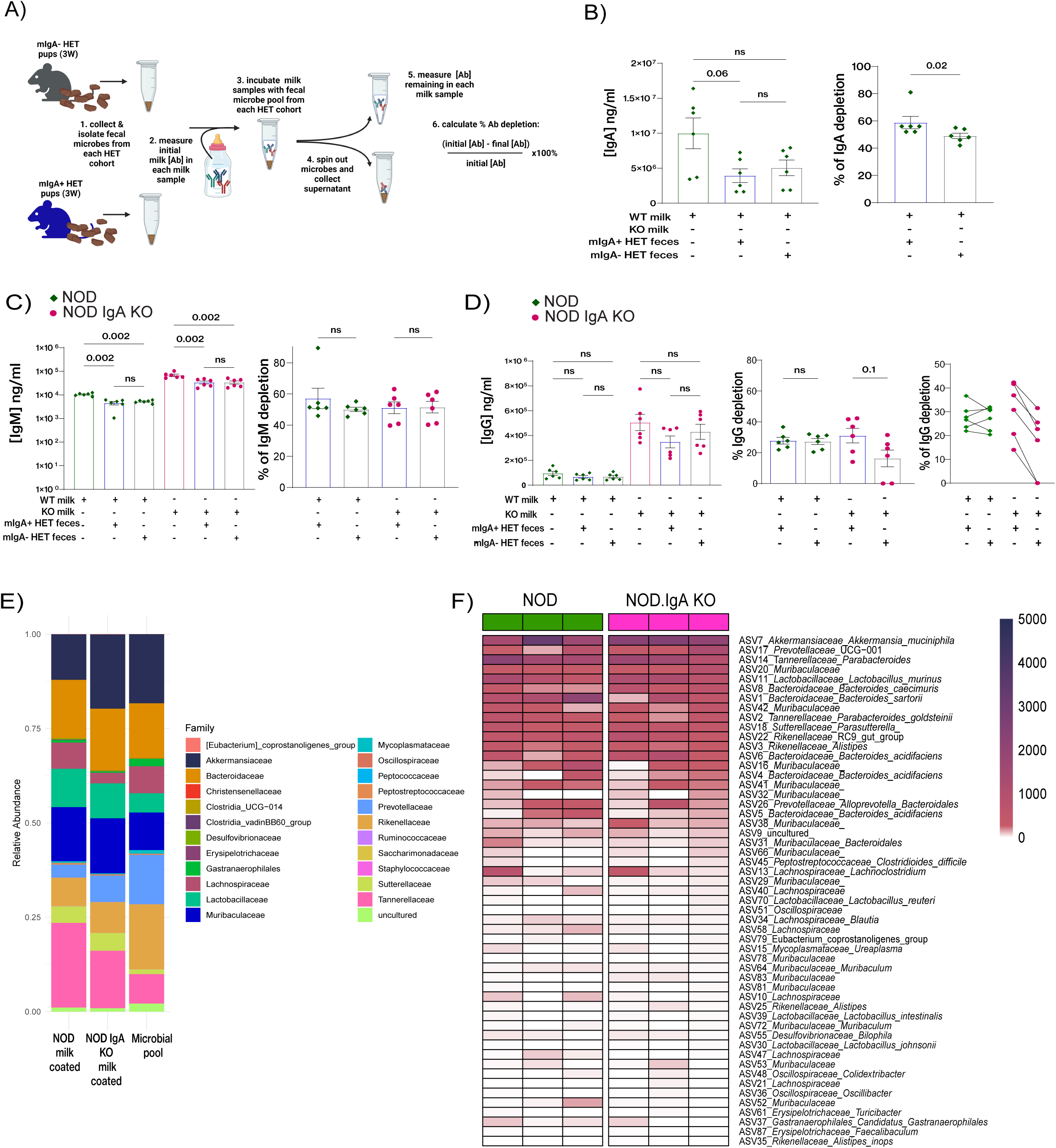
Milk antibodies from IgA-sufficient vs -deficient dams show differing potential to modulate offspring microbiota. (A) Schematic diagram of antibody depletion assays used to determine fecal microbe-binding capacity of milk antibodies from NOD and NOD.IgA KO dams. (B) Capacity of milk IgA, (C) IgM, and (D) IgG to coat fecal microbes from mIgA+ and mIgA-HET 3 week old pups. Bar graphs on the left of each panel indicate antibody concentrations before and after incubation with fecal microbes from each of the NOD.IgA HET pup cohorts, while bar graphs on the right indicate the % decrease for each milk antibody isotype after incubation with fecal microbes. The right most graph in panel (D) indicates the % depletion for each milk sample following incubation with mIgA+ HET or mIgA- HET fecal sample, where dots connected by a line indicate the same milk sample used. (E) 16S rRNA gene sequencing data indicating the relative proportions of bacterial families present in milk antibody-positive fractions of a fecal microbial pool after incubation with NOD milk (left column, n=8) or NOD.IgA KO milk (middle column, n=8) compared against the starting fecal bacterial pool derived from 3 week old pups from both HET cohorts (right column). (F) 16S sequencing data displayed as a heatmap comparing propensity of NOD vs NOD.IgA KO milk antibodies to coat a fecal bacterial pool. *For panels (B)-(D), data is shown as means ± SEM. p-values were determined using the Mann-Whitney U test. Each data point represents one mouse*.

IgM is also present in breast milk and can carry out similar functions to IgA in the gut lumen. Thus, we asked whether IgM from IgA-deficient milk might exhibit increased microbe coating abilities. However, we found that across all combinations of milk and fecal microbe pools, milk IgM showed approximately 50% depletion after incubation, with no enhanced coating by NOD.IgA KO milk IgM (FIGURE 10C). Surprisingly, IgG in NOD.IgA KO milk, which was 5- fold more abundant than NOD milk, was largely inadequate at coating fecal microbes from mIgA- HET pups (FIGURE 10D), further supporting our hypothesis that microbial communities in this offspring cohort were escaping modulation by milk antibodies. Furthermore, NOD.IgA KO milk IgG showed amplified coating of mIgA+ HET microbes, possibly indicating the presence of more immunogenic or potentially pathogenic microbial species in this cohort. Taken together, our results indicate milk IgG does not play a predominant role in microbiome modulation in the neonatal gut, raising questions regarding the antigenic repertoire of this antibody isotype and its primary role in neonatal development.

To identify which species were being recognized by milk antibodies, we employed combined IgA/IgG/IgM sequencing, where pooled fecal microbes from 3 week old mIgA+ HET and mIgA- HET mice that were coated by NOD and NOD.IgA KO milk antibodies were subjected to 16S rRNA gene amplicon sequencing. NOD milk displayed a capacity to recognize a broader range of bacterial species, likely due to the presence of IgA and a greater antibody content overall. Considering the lack of IgA in milk from IgA-deficient dams, we were surprised at how similar the milk antibody-coated microbial fractions looked (FIGURE 10E). Both milk types showed substantial ability to coat *Lactobacillaceae*, *Muribaculaceae*, *Bacteroidaceae*, and *Sutterellaceae* members, such that the coated microbe fractions showed enrichment of these taxa compared to the original fecal microbial pool. *Tannerellaceae* species were also substantially coated by both milk types, but to a higher degree by NOD milk. NOD milk also harbored more *Lachnospiraceae*-reactive antibodies, while NOD.IgA KO milk contained more antibodies recognizing *Prevotellaceae* and *Akkermansiaceae.* Notably, milk from IgA-deficient dams contained more *A. muciniphila*-reactive antibodies than did NOD milk (FIGURE 10F), providing a plausible mechanism for the reduced abundance of this species in offspring of IgA-deficient dams. Moreover, since 16S rRNA gene sequencing provides information on the relative abundance of microorganisms present, this data provides no information on the absolute abundance of bacteria coated by each milk sample, and by extension, on the fraction of bacteria escaping modulation.

Taken together, our data provide evidence for differential microbe-reactivities in the milk antibody repertoire between NOD and NOD.IgA KO dams, resulting in unique microbiome modulation in mIgA+ vs mIgA- HET offspring. Moreover, the data support IgA as the predominantly active antibody in breast milk for modulation of the microbiota, and a loss of this isotype leaves a significant proportion of the microbial community unsupervised.

## Discussion

The role of IgA in mucosal immunity and maintenance of GI homeostasis is well established. Microbes with pathogenic potential potentiate an IgA response in the GI tract, among other mucosal defense mechanisms, to preserve homeostasis. IgA deficiency causes intestinal dysbiosis, which has been shown to exacerbate obesity-related inflammation and insulin resistance^46,47^, and autoimmune central nervous system inflammation^48,49^. T1D patients also exhibit abnormal IgA responses to gut bacteria^60^, and several studies have reported a higher incidence of selective IgA deficiency in individuals with T1D^52–54^. The apparent disruption of GI homeostasis in individuals affected by T1D, particularly during the early pre-diabetic stages, suggests that compromised mucosal defenses could play a role in T1D pathogenesis. These observations collectively raise the critical question of whether IgA influences early diabetogenic events, either by promoting or preventing them.

In NOD mice, decades of studies illustrated the importance of the microbiome in T1D pathogenesis, highlighted by studies where genetic approaches that modified immunity altered the gut microbiome and led to significant changes in the manifestation of T1D^83–87^. Unexpectedly, we discovered that the loss of IgA in the NOD model did not elevate T1D incidence. Instead, we observed a dominant protective effect linked to maternal IgA deficiency, partly driven by alterations in the maternal microbiome and humoral immune factors present in the milk. Postnatal exposure to these factors influenced the development of the offspring’s microbiome, gastrointestinal mucosal immunity, and resilience, ultimately providing long-lasting protection against the disease.

A robust weaning reaction with appropriate GI colonization is critical in preventing pathological imprinting^22^. Here, we provide strong evidence that even when pathological imprinting has been written into one’s DNA, as is the case for both NOD mice and familial T1D in humans, a robust weaning reaction has the capacity to ameliorate its manifestation. Furthermore, where previous studies have evaluated the weaning reaction in black-or-white terms - comparing the presence or absence of GI colonization, we report that physiologically-relevant, subtle perturbations of the microbiome elicited by maternal cues have the capacity to bolster the weaning reaction and alter disease development. We investigated microbiome establishment and modulation in offspring, along with early host-mediated immune responses, with an interest in how these factors affected the GI environment, shaped early and long-term immunity, and influenced T1D susceptibility within a genetically-conducive environment. We observed that mIgA- HET pups exhibited a heightened GI inflammation post-weaning yet a reduced T1D incidence vs mIgA+ HET controls. We also observed an increased expansion of colonic iTreg, and increased GI barrier function later in life. Taken together, we propose a model where maternal dysbiosis induced by deficient immunity fosters a neonatal microbiome that is favorable for tolerance induction, and that supports GI imprinting to bolster intestinal barrier function. Importantly, our findings uncover a neonatal window of opportunity for the prevention of immuno-dysregulatory diseases such as T1D.

One consequence of exposure to IgA-deficient milk is the upregulation of humoral immunity in the pups, as reflected by higher numbers of activated CD4^+^ T cells in the bowel and MLN, increased proportions of MLN-residing Tfh cells and germinal center B cells, and heightened IgA production in the pups. Published studies^25^ have shown the ability of maternal antibodies, particularly IgG and IgA, to reduce Tfh numbers and suppress the development of germinal centers in the MLN. Our observation that newly weaned mIgA- HET pups exhibited increased germinal center activity may suggest a weakening of this maternal suppressive function in the absence of breast milk IgA. Aside from its role in facilitating antibody class-switching, Tfh can give rise to autoregulatory Tr1-like cells^88–90^. Whether these suppressive populations are altered by early life signals is an intriguing future avenue to pursue.

That IgA-deficient pups are refractory to maternally afforded T1D resistance suggests that induction of endogenous IgA is a critical prerequisite for such protection (supplemental FIGURE 1). The precise triggers that are required for effective tolerance induction are currently under investigation in our laboratory. Proteomic analysis of breast milk revealed considerable differences in immunomodulatory factors between milk from NOD vs. NOD.IgA KO dams at 10-12 days postpartum, with notable changes in the milk antibody isotype usage and antigen-binding repertoire. While this is reflective of maternal microbiome changes^91^, whether the antibodies are protective, and dependent on self vs foreign antigen recognition, are important future questions to answer. Breast milk antibodies have been shown to dampen intestinal CD4^+^ T lymphocyte activation and germinal center development during the weaning/early post-weaning period^19,25^, and strategies that target antibody signaling and transport will help tease apart underlying mechanisms.

Another means through which maternal antibodies could modulate offspring humoral immunity is through shaping the offspring microbiome. Sequestration of microbial antigens by maternal IgA in the gut lumen of offspring was the purported mechanism of maintained neonatal CD4^+^ T cell naivety^19^. Data from our milk antibody depletion assays support this theory, where we saw no increased capacity for microbe coating by IgM or IgG in IgA-deficient milk, thus leaving a significant proportion of mIgA-HET microbes unsequestered. 16S sequencing of milk antibody-coated fecal microbes indicated that several species in mIgA- HET pups may have escaped modulation by milk antibodies, while others were restrained to a greater degree. Furthermore, differential coating of microbiota communities could explain our observation that NOD.IgA HET offspring cohorts harbored enriched bacterial species that were not enriched in one dam cohort over the other.

Alternative mechanisms of maternal influence via breast milk include offspring microbiota modulation by non-antibody milk immune factors, differential microbiome seeding via introduction of breast milk-resident microorganisms, or microbiome-independent pathways of neonatal immune education via breast milk immuno-modulatory factors. One particularly attractive theory is that differences in IgG content and/or antigenic specificity may allow for introduction of unique microbial antigens to their respective progeny cohort via antibody-antigen complexes, thus influencing the establishment of tolerance to different antigens and expanding the lymphocyte repertoire. Support for this can be found in a number of animal studies employing neonatal introduction of foreign antigen, where tolerance and lymphocyte repertoire shaping results^77–79^. Our finding that milk from NOD.IgA KO dams produced heightened coating of *A. muciniphila* and *B. caecimuris*, likely via milk IgG, raises the possibility that increased early exposure to antigen from these species resulted in tolerance towards potential microbial mimitopes in mIgA- HET pups, and provides an attractive direction for further exploration.

Human studies of familial T1D have demonstrated that despite a vast diversity in parental genetic makeup, children consistently show a reduced risk of developing the disease when born to an affected mother over an affected father. Our findings recapitulate this inverse relationship, where maternal dysbiosis associated with IgA deficiency is protective against offspring T1D development. Our study highlights the existence and importance of a maternally afforded dominantly protective mechanism that can be leveraged to prevent T1D. Importantly, our data point to the strength and persistence of maternal effects that are sustained throughout different life stages, via gut imprinting and induction of tolerogenic mechanisms. Whether maternal signals also impact pancreatic islet development, function, and resilience in this scenario^92^ warrants further investigation.

Last but not least, in carrying out the above study, we strictly controlled for maternal effects, littermate effects, and cage sharing effects, which all contributed to the microbiome, immune and T1D modulation in our dam vs pup cohorts. Indeed, vertical transmission of the maternal microbiome has been shown to dominate over genetic effects as a driving force when PRR-deficient and -sufficient littermates shared the same microbiome^93^. We therefore advocate for careful examination of maternal influences in genetic models of disease, as a source of valuable yet often untapped mechanistic insights.

## Acknowledgments

We thank Dr. Margaret Conner, Scientific Director of the Gnotobiotics Core at Baylor College of Medicine in Houston, Texas, for the generous donation of C57BL/6 IgA deficient mice. We thank The Metabolomics Innovation Centre (Dr. Liang Li, Dr. Shuang Zhao, Dr. Xian Luo, and Xiaohang Wang) for their help with performing and analyzing metabolite data. We thank Dr. Aja Rieger and her team (Faculty of Medicine & Dentistry Flow Cytometry Facility) and Dr. Kacie Norton (Faculty of Science Advanced Microscopy Facility) at the University of Alberta for their help with this project. Funding for this project was provided by Canadian Institute of Health Research (CIHR), the Alberta Diabetes Institute (ADI), the University of Alberta, Li Ka Shing Institute of Virology (LKSIoV) and the Praespero Autoimmunity Fund. This work was supported in part by infrastructure CFI-JELF awards (O.J. 37833 and 39051), by the Canada Foundation for Innovation through SPP-ARC (Striving for Pandemic Preparedness - The Alberta Research Consortium). ES is supported by funding from the LKSIoV, University of Alberta and CIHR. LS is supported by LKSIoV, University of Alberta and ADI. ASC is funded by an NSERC-CREATE postdoctoral fellowship. ST is a Tier II Canada Research Chair in Immunometabolism and Diabetes.

## Author contributions

ES led the study, designed and conducted experiments, generated figures and wrote the manuscript. LS designed and conducted experiments, and generated figures. TJ analyzed 16S sequencing data, generated figures and advised on microbiome experimental design and data interpretation. ASC analyzed and generated figures associated with transcriptomic, proteomic, metabolomic and 16S sequencing data. HW and OJ performed proteomic analysis and analyzed data. MA provided technical support for experiments. PJB aided with interpretation of data and edited the manuscript. CH edited the manuscript. BW performed 16S sequencing and supervised microbiome experimental designs, analysis and interpretation. XCC supervised data interpretation and edited the manuscript. ST conceived and supervised the study and edited the manuscript.

## Methods

### Generation of the IgA-deficient NOD strain (NOD.IgA KO)

Male IgA deficient C57BL/6 mice, graciously donated by Dr. Margaret Conner, Scientific Director of the Gnotobiotics Core at Baylor College of Medicine in Houston, Texas, were mated with female NOD mice to produce progeny with one immunoglobulin heavy chain (IgH) gene bearing a deletion encompassing the complete ⍺ switch region and a portion of the ⍺ constant region^94^. Animals of the F1 generation were backcrossed to the NOD strain for 10 generations before mating two heterozygous animals to produce fully IgA-deficient NOD mice, denoted as NOD.IgA KO. Male and female NOD.IgA KO mice were then mated for >3 generations to produce multi-generational IgA-deficient progeny.

### Mating, weaning & fostering

NOD.IgA KO mice were generated via mating of NOD.IgA KO males and females. To generate NOD.IgA HET from IgA sufficient dams (mIgA+ HET), NOD females were mated with NOD.IgA KO males. NOD.IgA HET from IgA-deficient dams (mIgA- HET) were generated by mating NOD.IgA KO females with NOD males. All breeders were >10 weeks of age. Prior to mating, all mice were tested for glucosuria. Breeders that tested positive for glucosuria at any point during mating were retired. Litters arising from dams positive for glucosuria either during pregnancy or prior to weaning were not used for experiments. Within 24 hours of birth, sires were removed from the breeding cages in an effort to minimize paternal factor contribution. All pups were weaned at 21-22 days of age.

For fostering experiments, mIgA+ HET were fostered to IgA-deficient dams by dam exchange within 24 hours of birth. Fostered offspring were weaned at 21-22 days of age and followed for disease for 30 weeks.

All mice were bred and housed in specific pathogen free (SPF) conditions at 22°C with 12 hour light/dark cycle. Mice were housed with others of the same genotype and maternal lineage. This project (AUP00003665) received research ethics approval from the University of Alberta Research Ethics Board, and all animal use and experimentation was conducted with approval from relevant Animal Care and Use Committees (ACUC) and met Canadian Council on Animal Care (CCAC) guidelines.

### Disease monitoring

NOD, NOD.IgA HET & KO female mice were monitored weekly for glucosuria using Diastix (Ascensia Diabetes Care). A urine glucose concentration >0.5 g/dL constituted a positive test, whereupon mice were euthanized. Mice were followed until 30 weeks of age.

### Collection and processing of tissue and blood

At indicated age, mice were euthanized in a CO_2_ chamber and relevant tissues and organs removed for processing. MLN and PLN were placed in cold PBS, then manually homogenized and passed through a 70 μm filter to create a single cell suspension. Cells were washed in cold PBS, centrifuged at 1500 rpm for 5 min, resuspended in RPMI and stored on ice for cell counting. Large bowels were excised and measured for length, followed by manual removal of fecal contents and mucus. They were weighed, cut longitudinally, sectioned into ∼5 mm segments and stored in PBS on ice. A 10 cm section of the small bowel adjacent to the cecum was excised, relieved of its mucus layer and sectioned similar to the large bowel. Bowel segments were incubated in 10 ml stripping buffer (HBSS + 2% FBS, 5mM EDTA, 15mM HEPES) and incubated at 37°C with shaking for 20 min (x2), rinsed in 10 ml of ice cold wash buffer (HBSS + 2% FBS, 15mM HEPES), minced with scissors to create a pulp, transferred into 10 ml digestion buffer (RPMI + 10%FBS, Penn/Strep, L-glutamine, 1 mM NaPyruvate, β-ME, 50 U/ml DNase, 0.5 mg/ml Collagenase IV), incubated at 37°C for 45 min with shaking, passed through a 70 μm filter, rinsed with cold PBS, centrifuged at 1800 rpm for 5 min, resuspended in RPMI and cells were counted.

For each tissue, approximately 1×10^6^ cells were transferred to a 96 well plate and centrifuged at 1800 rpm for 5 min. The supernatant was removed and cells were blocked with Fc-block in FACS buffer (1/100 dilution) for 10 min on ice. An additional aliquot of FACS buffer containing antibodies (1/250 dilution) directed against relevant extracellular markers was added, and incubated at 4°C for 30 min. Cells were washed, centrifuged at 1800 rpm, resuspended in FoxP3 fix/perm buffer (eBiosciences), and incubated at 4°C for 45 min. Cells were centrifuged and resuspended in kit Fixation buffer and left overnight at 4°C. The next morning, cells were centrifuged and resuspended in kit Fixation buffer containing antibodies (1/250 dilution) directed against relevant intracellular targets. Samples were incubated at 4°C for 30 min, washed, centrifuged, and resuspended in Fixation buffer, then analyzed by flow cytometry on a BD LSRFortessa X-20. FlowJo (BD, v10.8) was used to analyze all cytometry data. Information on the antibodies used can be found in the Key Resources Table.

Blood was collected within 5 min of euthanization using a 23 gauge needle and drawing directly from the heart. The needle was then removed from the syringe before expelling the blood into a 1.5 ml Eppendorf tube. Blood was kept on ice for 1hr, then centrifuged at 10,000g for 10 min. Serum was carefully drawn off the top without disturbing the coagulated blood cell pellet and transferred to a fresh 1.5 ml Eppendorf tube for storage at −20°C.

### ELISA

Fecal samples collected from female mice at 5 weeks of age were assayed for immunoglobulin and Lcn-2 content using ELISA kits (Mabtech; R&D Systems). Protein was extracted from fecal samples by manual homogenization of 2-3 pellets in PBS + 0.05% Tween20 followed by 4 hr of agitation at 4°C, centrifugation (10,000g 10min @ 4°C), and collection of supernatant. Total protein concentration was determined using Nanodrop protein quantification (Fisher Scientific). Fecal samples were diluted 1/10^2^ (IgA), 1/10 (IgG & IgM), and 1/10 – 1/10^2^ (Lcn-2). Serum was diluted 1/10^3^ (IgA) and 1/10^5^ (IgG). Breast milk was diluted 1/10^3^ – 1/10^4^ (IgA), 1/10^3^ – 1/10^4^ (IgG), and 1/10^4^ – 1/10^5^ (IgM). All ELISA were run in duplicate or triplicate.

### Gut Permeability Assays

At 7, 10 and 15 weeks of age, female mice were assessed for gut permeability via oral gavage with FITC-dextran beads (FD4; 0.4mg/g body weight; Cedarlane). Blood (approx. 80μl) was collected from the saphenous vein 4 hrs post-gavage, centrifuged at 10,000g for 10 min and the serum assayed for FD4 concentration against a standard curve.

### Cytokine/receptor expression analysis (qPCR)

Ileum sections were examined for expression of various mRNA. After euthanasia via CO_2_ inhalation and removal of bowel contents, 2 cm section of small bowel tissue was excised approximately 1 cm from the cecum and frozen at −80°C in Trizol reagent (Invitrogen). To extract RNA, samples were thawed on ice, homogenized manually, and topped up to 1 ml with Trizol reagent. 200 μl chloroform was added, samples were shaken vigorously, allowed to separate at room temperature for 5 minutes, then spun at 12,000g for 15 min at 4°C. The top aqueous layer (approx 500 μl) was carefully removed and transferred to a fresh collection tube containing equivalent volume of cold isopropanol. Samples were inverted to mix, incubated on ice for 10 min, then applied to an RNeasy spin column (Qiagen) for RNA purification following the Qiagen RNeasy kit protocol. To elute RNA, 40-75 μl of RNase/DNase-free H_2_O was applied to the column,incubated at room temperature for 1 min, then spun for 1 min at 10,000g. RNA was quantified using OD 260 reading on a Nanodrop spectrophotometer (ThermoFisher) and purity assessed using the 260/280 and 260/230 ratios. RNA samples were stored at −80°C.

cDNA was generated from purified RNA samples using the Superscript III Reverse Transcription kit (Invitrogen) and used as template for quantitative PCR reactions of various genes of interest. Primer sequences were either designed and optimized with both Advance (Wisent) and PerfeCTa Supermix (Quantabio) SYBR green qPCR reaction mixes. qPCR reactions were performed on CFX96 TouchReal-Time PCR Detection System (BioRAD) and carried out in either duplicate or triplicate.

Purified RNA was sent to Novogene Corporation Inc. for bulk mRNA sequencing on Illumina NovaSeq X Plus platform. mRNA was purified from total RNA, cDNA synthesized using random hexamers, followed by end repairing and A-tailing, adapter ligation, size selection, PCR amplification, and purification. cDNA libraries were quantified and analyzed for size distribution. Following sequencing, raw data underwent quality control, including examination of error rate distribution, GC-content distribution and data filtering (removal of reads with adaptor contamination, N>10%, and >50% low quality base calls). See Supplemental Material for data quality summary.

### Histology & scoring of pancreata

Pancreata were excised from freshly euthanized 12-13 week female mice, placed into cassettes, immediately fixed in 10% formalin for >48 hr, then transferred to 70% ethanol for >24 hr. Samples were dehydrated and paraffin embedded by immersion in 50% ethanol (1 hr), 70% ethanol (1 hr), 80% ethanol (1 hr), 95% ethanol (1 hr), 100% ethanol (1.5 hr X3), toluene (1.5 hr X3), and paraffin wax at 60°C (2 hr X2).

Tissue was then removed from cassettes, placed in hot wax molds and allowed to harden. 5 μm sections were placed on glass slides and allowed to dry overnight at 37°C. Deparaffinization & rehydration was performed by immersion in toluene (5 min X2), 100% ethanol (2 min X2), 90% ethanol (2 min), 70% ethanol (2 min), 50% ethanol (2 min), and distilled H_2_O (2 min). Hematoxylin & eosin (H&E) staining was performed by immersion in hematoxylin (2 min) followed by rinsing in gently running H_2_O for 15 min, immersion in 70% ethanol (2 min), eosin (30 s), 100% ethanol (2 min X2), and toluene (2 min X2). Slides were coverslipped and sealed with DPX (distyrene + a plasticizer + xylene) mountant, then allowed to harden overnight at 37°C. For each pancreas, 8X 5 μm sections were analyzed with a distance of ∼60 μm between any two subsequent sections, allowing for analysis throughout the depth of the tissue. Islets were scored for degree of immune cell infiltration, where a score of 0 indicates no immune cells present, 1 indicates immune cells are visible on islet periphery, 2 indicates an infiltration <25% of islet area, 3 indicates an infiltration of 25% - 50% of islet area, and 4 indicates an infiltration >50% of islet area. Islet scores from all 8 sections were summed, and the average islet infiltration score was calculated and plotted using GraphPad Prism 9 (https://www.graphpad.com).

### Histology & scoring of bowel sections

Sections of large bowel approximately 3 cm in length were harvested from freshly euthanized female mice such that each section contained a complete fecal pellet. Whenever possible, a section from the middle of the large bowel was collected. The tissue was immediately placed in Carnoy’s solution (60% ethanol, 30% chloroform, 10% glacial acetic acid) and fixed overnight at 4°C. The tissue was removed from the Carnoy’s solution, cut transversely through the fecal pellet with a sharp blade, then transferred to 100% ethanol at 4°C. The 100% ethanol was changed twice over a span of 2-3 days, then the tissue was dehydrated and paraffin embedded as outlined above. The bowel sections were placed in the paraffin block such that sectioning of the tissue resulted in transverse sections. 7 μm sections were mounted on glass slides and dried overnight at 37°C.

Slide-mounted sections were deparaffinized and rehydrated as outlined above, then fluorescence *in situ* hybridization (FISH) was performed. Slides were incubated in wash buffer (0.9M NaCl, 20mM Tris-HCL, pH 7.2) at 50°C for 10 min, then incubated with a fluorescently labelled pan-bacterial 16S probe (Eub338, 5’GCTGCCTCCCGTAGGAGT3’; IDT) diluted to 100 nM in wash buffer + 0.1% SDS at 50°C for overnight in a dark humidification chamber. Slides were washed in 50°C wash buffer for 10 min, then for 8 min in fresh wash buffer at room temperature. Lectin from *Ulex Europaeus* (agglutinin; Invitrogen) diluted to 20 μg/ml in PBS was added to the slides and incubated for 2 hr at room temperature in a dark humidification chamber. Slides were washed for 8 min in wash buffer at room temperature, then mounted with ProLong™ Gold Antifade Mountant with DNA Stain DAPI (Invitrogen). FISH slides were visualized and images collected on a Cytation 10 multimode plate reader (Agilent Biotek) using the 4X magnification lens. ImageJ software (NIH) was used to assess the thickness and integrity of the mucosal layer in a blinded fashion. For thickness assessment, 6 areas were chosen at regular intervals around the tissue section and the mucosal layer measured at each. A minimum of 5 sections per sample were assessed, and an average thickness was generated and plotted using GraphPad Prism 9.

### Microbiota analysis

Fecal pellets were collected as they were excreted into sterile 1.5 ml Eppendorf tubes and stored immediately at −80°C. Frozen fecal pellets were weighted and DNA was extracted using the QIAamp Fast DNA Stool Mini Kit (Qiagen) following the protocol provided in the kit with inclusion of a 1 min bead-beating step with garnet beads (BioSpec Products) in a FastPrep-24 5G homogenizer (MP Biomedicals). Library prep was performed on purified DNA, targeting the V3-V4 region of the 16S rRNA gene according to the 16S metagenomic Sequencing Library Preparation protocol, and sequencing was performed on Illumina MiSeq v2 by McGill Genome Centre. Raw data sequences were processed using the Quantitative Insights Into Microbial Ecology 2 (QIIME2, v2021.2) pipeline, and downstream analyses were performed in R (https://www.r-project.org/).

### Metabolomic analysis

Fecal pellets were collected as they were excreted into sterile 1.5ml Eppendorf tubes. Samples were immediately stored at −80°C. Frozen fecal material was delivered on dry ice to The Metabolomics Innovation Center (TMIC) at the University of Alberta for metabolomic profiling using High Performance Chemical Isotope Labeling Metabolomics. Analysis of raw data was carried out by TMIC personnel and entered into Microsoft Excel spreadsheets (.xlsx & .csv files). The .xlsx and .csv files were uploaded to MetaboAnalyst 5.0 (https://www.metaboanalyst.ca/home.xhtml) for identification of enriched or depleted metabolites, pathway analysis and generation of figures.

### Breast milk collection & proteomic analysis

Milk was collected from breastfeeding dams 10-13 days postpartum. Dam and pups were physically separated for 4 hrs such that visual, olfactory and audible connections were maintained but pups could not nurse. The dam was then anesthetized with isoflurane and given oxytocin (2 IU/kg) intraperitoneally. Milk was expressed by gently massaging inguinal nipples and collecting milk droplets with a sterile pipet. Upon collection of approx. 80-100 μl of milk, the dam was allowed to recover and then returned to her litter. Breast milk was stored at −20°C.

Breast milk samples were thawed at 37°C for 5 min, then mixed thoroughly. Milk was diluted 10 fold in PBS + 0.28% SDS and transferred to a 0.2 ml PCR tube nested inside a 1.5 ml Eppendorf tube. The samples were centrifuged at 14,000 rpm for 10 min to separate the fat from the aqueous phase. The aqueous layer was collected and transferred to a fresh 0.2 ml tube and stored at −20°C.

Breast milk samples were prepared for mass spectrometry analysis using the ProTrapXG column (Allumiqs Corporation). Milk protein (∼20µg) for NOD and NOD IgA KO (n=3 each) was precipitated in a filtration cartridge with 400µL acetone for 30min at room temperature. Protein aggregates were collected by centrifugation at 2500g for 2 min, supernatant discarded at 400g for 5 min, and washed with 400µL acetone at 400g for 2 min. Collected proteins were resolubilized in 8M urea, vortexed, sonicated, and left at R.T for 30 min. Then, 100mM Tris (pH=8) was added, followed by reduction with 10mM DTT at 37°C for 30 min and alkylation with 20mM iodoacetamide for 45 min in dark. Trypsin (Promega) was added 50:1 (protein:trypsin) and incubated on a rotator overnight at R.T. Trypsin reaction was stopped by the addition of trifluoroacetic acid to a final concentration of 1% and incubated for 10 min at R.T. Peptide mixture was then centrifuged at 13000g for 10 min, with the supernatant collected for peptide clean-up and desalting on the ProTrapXG SPE cartridge by manufacturer recommendations. The tryptic peptides solution was dried by a centrifugal evaporator (Genevac EZ-2 4.0 series) and resuspended in 0.1% formic acid for LC-MS/MS analysis.

Peptide samples were analyzed via mass spectrometry using a nanoflow-HPLC (Thermo Scientific EASY-nLC 1200 system) coupled to an Orbitrap Fusion Lumos Tribrid Mass Spectrometer (Thermo Fisher Scientific). The peptide samples underwent reverse phase separation on an analytical column (Aurora Ultimate nanoflow UHPLC column 25 cm x 75 µm ID, 1.7 µm C18, 120 Å; IonOpticks inc.). They were eluted over a 120 min non-linear gradient from 0% to 80% acetonitrile in 0.1 % formic acid. Data-independent acquisition was conducted over 375 to 2000 m/z at a resolution of 120,000 with a maximum injection time of 50ms. Spectral data was analyzed on Spectronaut (v18.7.240506.55695) using factory settings against a UniProt database of the mouse proteome (Mus musculus Proteome ID UP000000589, download date 06/21/2021). Search parameters included Trypsin/P cleavage (max 2 missed cleavages), fixed modification of carbamidomethylation (C), variable modifications of oxidation (M) and deamidation (N/Q), and maximum variable modifications of 3. The false discovery rate was set to 1%, and precursor filtering was performed with a Q-value cutoff of 1%. Quantification was done at the MS2 level based on the peak area, with differential abundance comparisons done by unpaired *t-*tests using MS2 quantities. Significantly changed proteins were filtered by a fold change threshold of 1.5 and p-value less than 0.05.

### Breast milk antibody specificity assay

Breast milk samples were thawed at 37°C for 5 min, then mixed thoroughly. Aliquots from samples were diluted 1/10 in FACS buffer containing PE conjugated antibody (either IgA, IgG or IgM). Samples were incubated on ice for 1.5 hrs.

Meanwhile, fecal pellets were collected from NOD.IgA HET pups (25 days of age) into sterile PBS+10% glycerol. In an anaerobic chamber, fecal matter was pooled, homogenized and passed through a 40 μm filter, then rinsed with sterile PBS. Flowthrough was spun in a centrifuge at 14,000g for 3 min, then resuspended in 1% paraformaldehyde (PFA) for 45 min on ice for fixation to preserve obligate anaerobic bacteria. Fecal pellets were then pelleted again at 14,000g for 3 min and resuspended in FACS buffer. Fecal pellets from 14 week NOD females and SCID NOD females were similarly collected and processed for use as controls.

An aliquot of fecal slurry + FITC conjugated SYTO16 (BioLegend) were added to each milk sample, mixed and incubated on ice for 45 min. Microbes were washed with FACS buffer, spun at 14,000g for 3 min, and resuspended in sorting buffer (PBS + 0.5%BSA + 2mM EDTA). PE-nanobeads (BioLegend) were added to each sample and incubated on ice for 15 min. Microbes were then washed with sorting buffer, pelleted as above, and resuspended in fresh sorting buffer. Samples were placed in magnets (Miltenyi) for 5 min, then supernatant was poured off. This was repeated twice for a total of 3 positive selection steps. The positively selected microbes remaining in the tubes were washed with PBS, pelleted as above, and the microbial pellet stored at −80°C to await 16S rRNA gene sequencing. An aliquot of each sample was analyzed with flow cytometry to ascertain degree of enrichment of each fraction.

### Breast milk antibody depletion assay

Fresh fecal pellets were collected from mIgA+ HET and mIgA-HET pups at 3 weeks of age, homogenized and passed through a 40 μm filter, then rinsed with sterile PBS. Flow-through was centrifuged at 14,000g for 3 min, then fixed in 1% PFA for 45 min on ice. Fecal material was pelleted at 14,000g for 3 min and resuspended in FACS buffer. Fecal microbes were resuspended and incubated in breast milk diluted 1:10 for 1hr @ room temperature, pelleted again and the supernatant retained for [Ab] determination. Initial [Ab] for each milk sample was determined, and the % depletion of each (microbe + milk) combination was calculated by subtracting the final from the initial [Ab], dividing by the initial [Ab] and multiplying by 100.

---

**Table 1.** List of antibodies for flow cytometry.

| **Antibody (anti-mouse)** | **Clone** | **Supplier** | **Catalog No.** |
| --- | --- | --- | --- |
| CD45 BV711 | 30-F11 | BioLegend | 103147 |
| CD3 PE Cy7 | 17A2 | BioLegend | 100219 |
| CD19 APC Cy7 | 1D3/CD19 | BioLegend | 152411 |
| CD4 Alexa Fluor 488 | GK1.5 | BioLegend | 100425 |
| CD8 BV605 | 53-6.7 | BioLegend | 100743 |
| TCR$\gamma\delta$ APC | GL3 | BioLegend | 118115 |
| FOXP3 PE | QA20A67 | BioLegend | 118903 |
| RORγt PE eFl 610 | AFKJS-9 | eBiosciences | 61-6981-82 |
| CD90.2 Alexa Fluor 700 | 30-H12 | BioLegend | 105320 |
| NKp46 BV650 | 29A1.4 | BioLegend | 137635 |
| Aqua Zombie live/dead stain | N/A | BioLegend | 423101 |
| CD138 BV650 | 281-2 | BioLegend | 142517 |
| BLIMP1 APC | 5E7 | BioLegend | 150007 |
| IgA PE | mA-6E1 | Fisher Scientific | 12-4204-82 |
| CD95 BV605 | SA367H8 | BioLegend | 152612 |
| GL7 Alexa Fluor 488 | GL7 | BioLegend | 144612 |
| CD86 PE Dazzle | GL-1 | BioLegend | 105041 |
| B220 Alexa Fluor 700 | RA3-6B2 | BioLegend | 103232 |
| CXCR5 BV711 | L138D7 | BioLegend | 145529 |
| PD1 BV605 | 29F.1A12 | BioLegend | 135219 |
| SIGLEC H PE | 551 | BioLegend | 129605 |
| CD44 APC | QA19A43 | BioLegend | 163603 |
| IgM PE | RMM-1 | BioLegend | 406508 |
| IgG PE | Poly4053 | BioLegend | 405307 |
| SYTO 16 | N/A | BioLegend | S7578 |
| TCRβ PE Cy5 | H57-597 | BioLegend | 109210 |

**Table 2.** List of reagents.

| **Item** | **Catalogue No.** | **Supplier** |
| --- | --- | --- |
| Diastix | 2804B | Ascensia Diabetes Care |
| Hank's Balanced Salt Solution (HBSS) | 311-513-CL | Wisent |
| EDTA | AC118445000 | Thermo Fisher Scientific |
| HEPES | ICN1688449 | MP Biomedicals Inc |
| RPMI 1640 medium | 350-002-CL | Wisent |
| Fetal bovine serum | F1051 | Sigma |
| Penn/Strep | P4333 | Sigma |
| L-glutamine | 25030081 | Gibco |
| NaPyruvate | 11-360-070 | Thermo Fisher Scientific |
| β-Mercaptoethanol | 60-24-2 | Sigma |
| MEM Non-essential amino acids | 11140-050 | Thermo Fisher Scientific |
| Collagenase IV | 17104019 | Gibco |
| DNase | D5025 | Sigma |
| Fc block | 422301 | BioLegend |
| Phosphate buffered saline (PBS) | BP399-20 | Thermo Fisher Scientific |
| Tween20 | MP1TWEEN201 | MP Biomedicals Inc |
| FITC-dextran beads | FD4-250MG | Cedarlane |
| isopropanol | A4641 | Thermo Fisher Scientific |
| Trizol | 10296010 | Invitrogen |
| Advance SYBR green qPCR reaction mix | 801-001-AR | Wisent |
| PerfeCTa SYBR green Supermix qPCR reaction mix | 95054-500 | Quantabio/VWR |
| 10% formalin | 22-050-104 | Thermo Fisher Scientific |
| toluene | T324-1 | Thermo Fisher Scientific |
| paraffin wax | 18614673 | Thermo Fisher Scientific |
| hematoxylin | H345-25 | Thermo Fisher Scientific |
| eosin | LC140258 | Thermo Fisher Scientific |
| DPX | 50980369 | Electron Microscopy Sciences |
| agglutinin | L32476 | Invitrogen |
| ProLong™ Gold Antifade Mountant | P36931 | Invitrogen |
| garnet beads, 20mm | 11079120gar | BioSpec Products |
| SYTO16-FITC | S7578 | BioLegend |
| Bovine Serum Albumin | A8806 | Sigma |
| PE-nanobeads | 480080 | BioLegend |
| vancomycin | 111002498 | MilliporeSigma |
| neomycin | AAJ6149914 | Thermo Fisher Scientific |
| ampicillin | BP1760-5 | Thermo Fisher Scientific |
| metronidazole | AC210345000 | Thermo Fisher Scientific |
| amphotericin B | SV3007801 | Thermo Fisher Scientific |
| the ProTrapXG column |  | Allumiqs Corporation |
| acetone | 423240040 | Thermo Fisher Scientific |
| urea | AM9902 | Invitrogen |
| Tris | 17926 | Thermo Fisher Scientific |
| Dithiothreitol (DTT) | D1532 | Invitrogen |
| iodoacetamide | 122275000 | Thermo Fisher Scientific |
| trypsin | V5113 | Promega |
| trifluoroacetic acid | 432295000 | Thermo Fisher Scientific |
| 0.1% formic acid | 270480250 | Thermo Fisher Scientific |
| Aurora Ultimate nanoflow UHPLC column 25<br>cm x 75 µm ID, 1.7 µm C18, 120 Å | AUR3-25075C18 | IonOpticks Inc. |
| acetonitrile | 047138-M1 | Thermo Fisher Scientific |

**Table 3.** List of commercial kits.

| **Kit Name** | **Catalog No.** | **Supplier** |
| --- | --- | --- |
| FoxP3/Transcription Factor Staining Buffer Set | 00-5523-00 | eBioscience |
| ELISA Flex: Mouse IgA (ALP) | 3865-1AD-6 | Mabtech, Inc |
| Mouse IgM ELISA BASIC kit (HRP) | 3885-1H-6 | Mabtech, Inc |
| Mouse IgG ELISA BASIC kit (ALP) | 3825-1AD-6 | Mabtech, Inc |
| Mouse Lcn-2/NGAL DuoSet ELISA | DY1857-05 | R&D Systems |
| Superscript III Reverse Transcription kit | 4368814 | Invitrogen |
| QIAamp Fast DNA Stool Mini Kit | 51604 | Qiagen |
| RNeasy micro kit (50) | 74004 | Qiagen |

**Table 4.** List of hardware & software.

| **Hardware/Software** | **Supplier** |
| --- | --- |
| FlowJo (BD, v10.8) | BD Biosciences |
| GraphPad Prism 9.0.1 | GraphPad Software, LLC |
| Cytation 10 multimode plate reader | Agilent Biotek |
| CFX96 Touch Real-Time PCR Detection System | BioRAD |
| NanoDrop 1000 | Thermo Fisher Scientific |
| ImageJ | National Institutes of Health (NIH) |
| FastPrep-24 5G homogenizer | MP Biomedicals |
| MetaboAnalyst 5.0 | https://www.metaboanalyst.ca/home.xhtml |
| Enrichr | https://maayanlab.cloud/Enrichr/ |
| BD LSRFortessa X-20 | BD Biosciences |
| QIIME2, v2021.2 | https://docs.qiime2.org/ |
| R | https://www.r-project.org/ |
| Thermo Scientific EASY-nLC 1200 system | Thermo Scientific |
| Orbitrap Fusion Lumos Tribrid Mass Spectrometer | Thermo Scientific |
| Spectronaut® (v18.7.240506.55695) | Biognosys |

