## Supplemental Figures for "Maternal humoral factors modulate offspring gut immune homeostasis to mitigate diabetes development"

a)

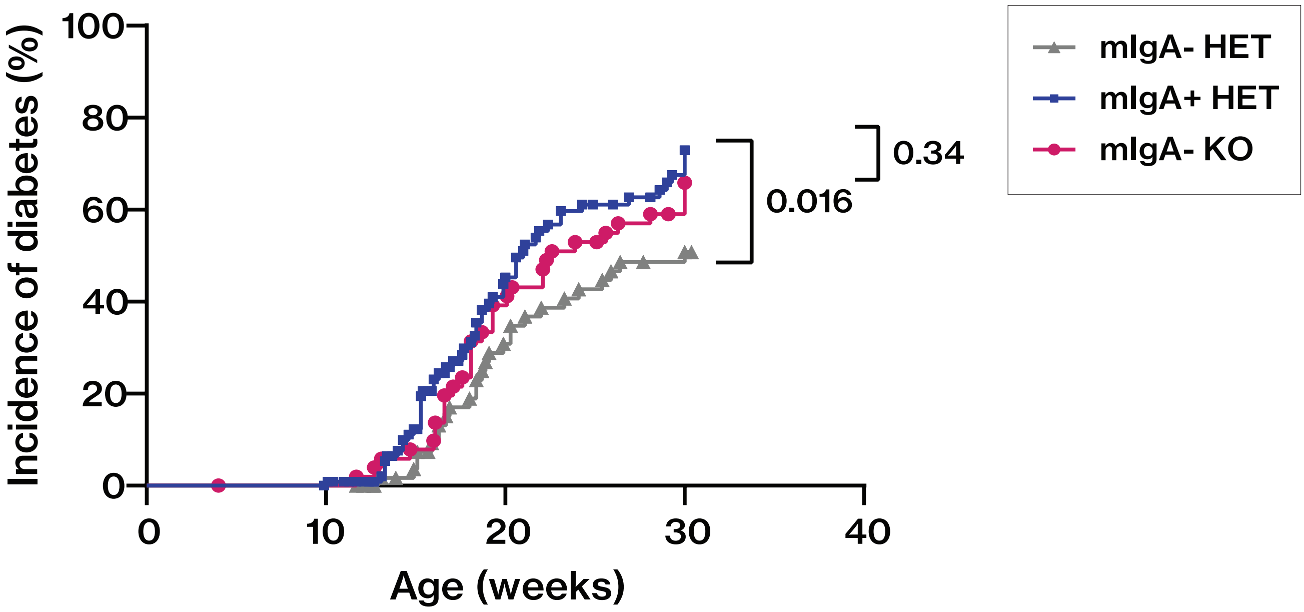


**Supplemental Figure 1. IgA-deficient offspring born to IgA-deficient dams do not show reduced T1D incidence**

(a) Incidence curve comparing T1D incidence in IgA-deficient female offspring born to IgA-deficient dams (mIgA- KO; pink circles), IgA-sufficient female offspring born to NOD dams (mIgA+ HET females; blue squares) and IgA-sufficient female offspring born to IgA-deficient dams (mIgA- HET females; grey triangles). *p-value was calculated using Log-rank test. Each data point represents one mouse.*

a)

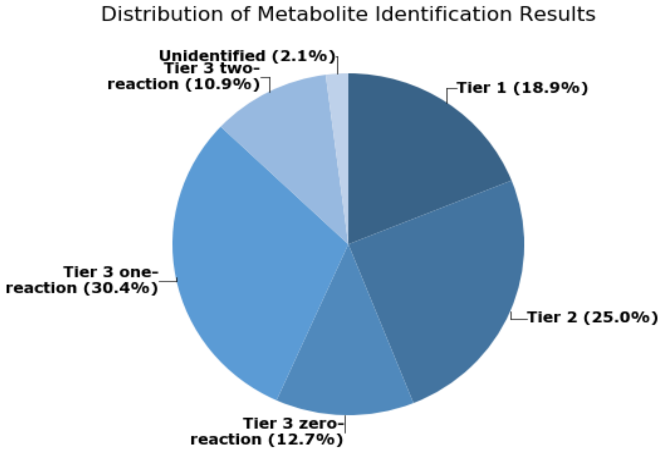


**Supplemental Figure 2. Detection and identification of fecal metabolites from NOD.IgA HET pups at 5 weeks of age**

(a) Collectively, 4422 metabolites were detected in fecal samples from mIgA+ HET (n=8) and mIgA- HET (n=9) pups at 5 weeks of age using high performance chemical isotope labeling LC-MS. 548 (18.9%) of these were identified using accurate mass and retention time, and searching against a labeled metabolite library (tier 1). An additional 918 (25.0%) metabolites were identified using accurate mass and predicted retention time, and searching against a linked identity library (tier 2).

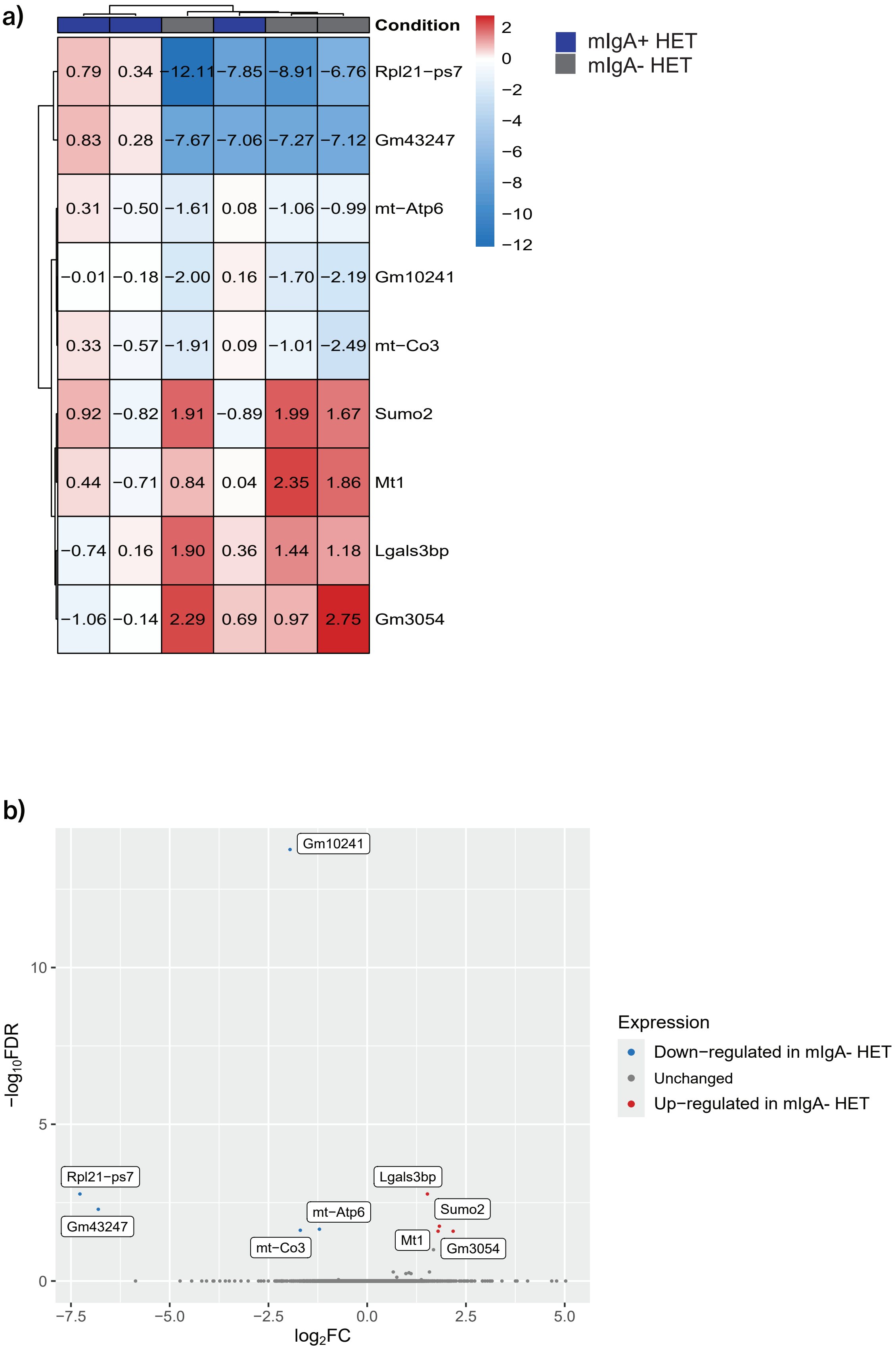


**Supplemental Figure 3. Transcriptomic analysis of colon tissue at 3 weeks reveals few differences between HET cohorts**

(a) Heatmap and (b) volcano plot showing differential expression of nine genes in colon tissue from 3 week old mIgA- vs mIgA+ HET pups.

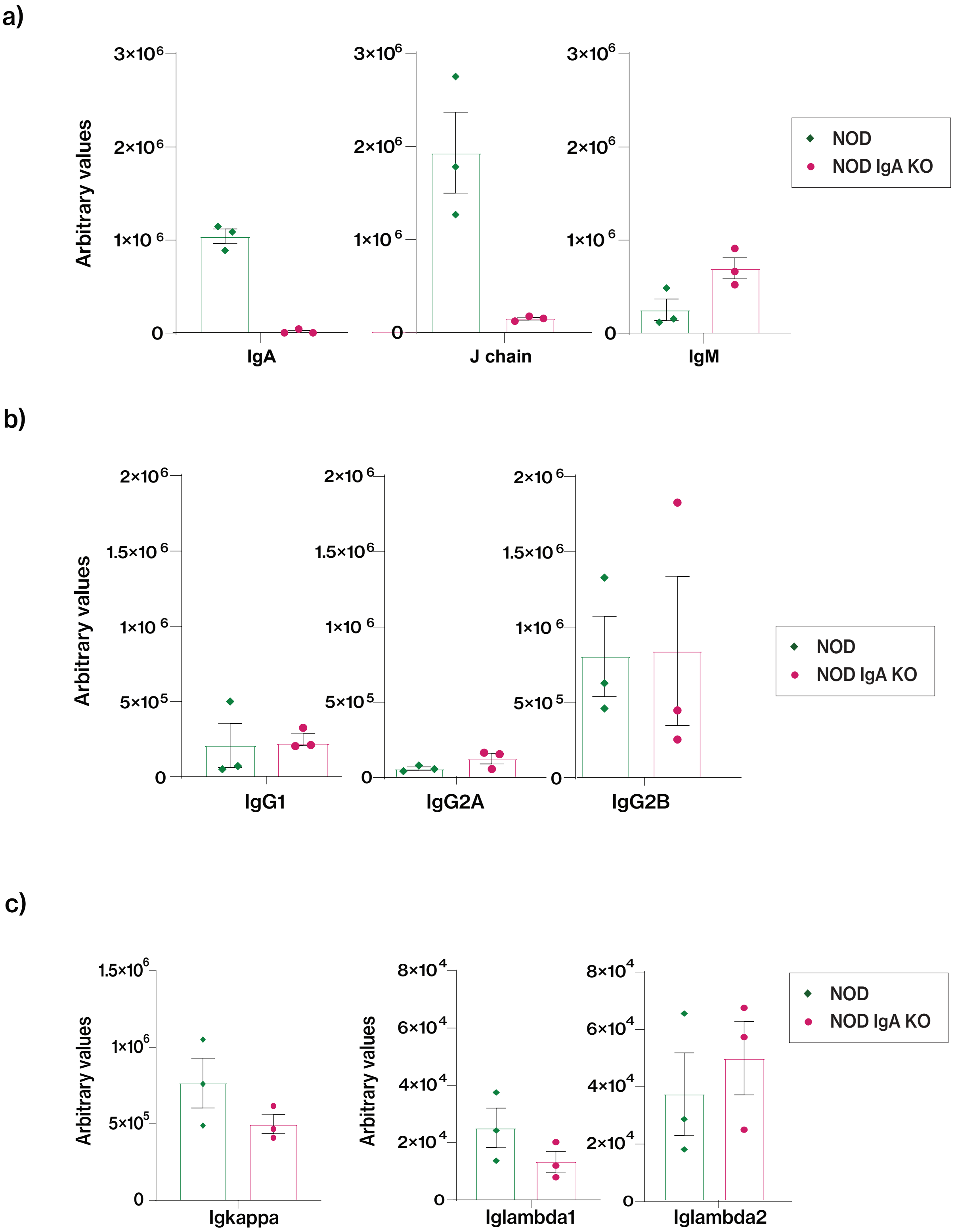


**Supplemental Figure 4. Comparison of milk immunoglobulin peptide abundances in IgA-sufficient vs. -deficient dams**

Analysis of milk protein content in NOD (green diamonds) and NOD.IgA KO milk (pink circles) collected at post-partum day 10-13, comparing relative abundances of (a) $\alpha$ constant region, J chain, and $\mu$ constant region peptides, (b) $\gamma$1, $\gamma$2A & $\gamma$2B constant region peptides, and (c) $\kappa$ constant region, $\lambda$1 constant region and $\lambda$2 constant region peptides. *p-values were determined using Mann-Whitney U test. Each data point represents one mouse.*

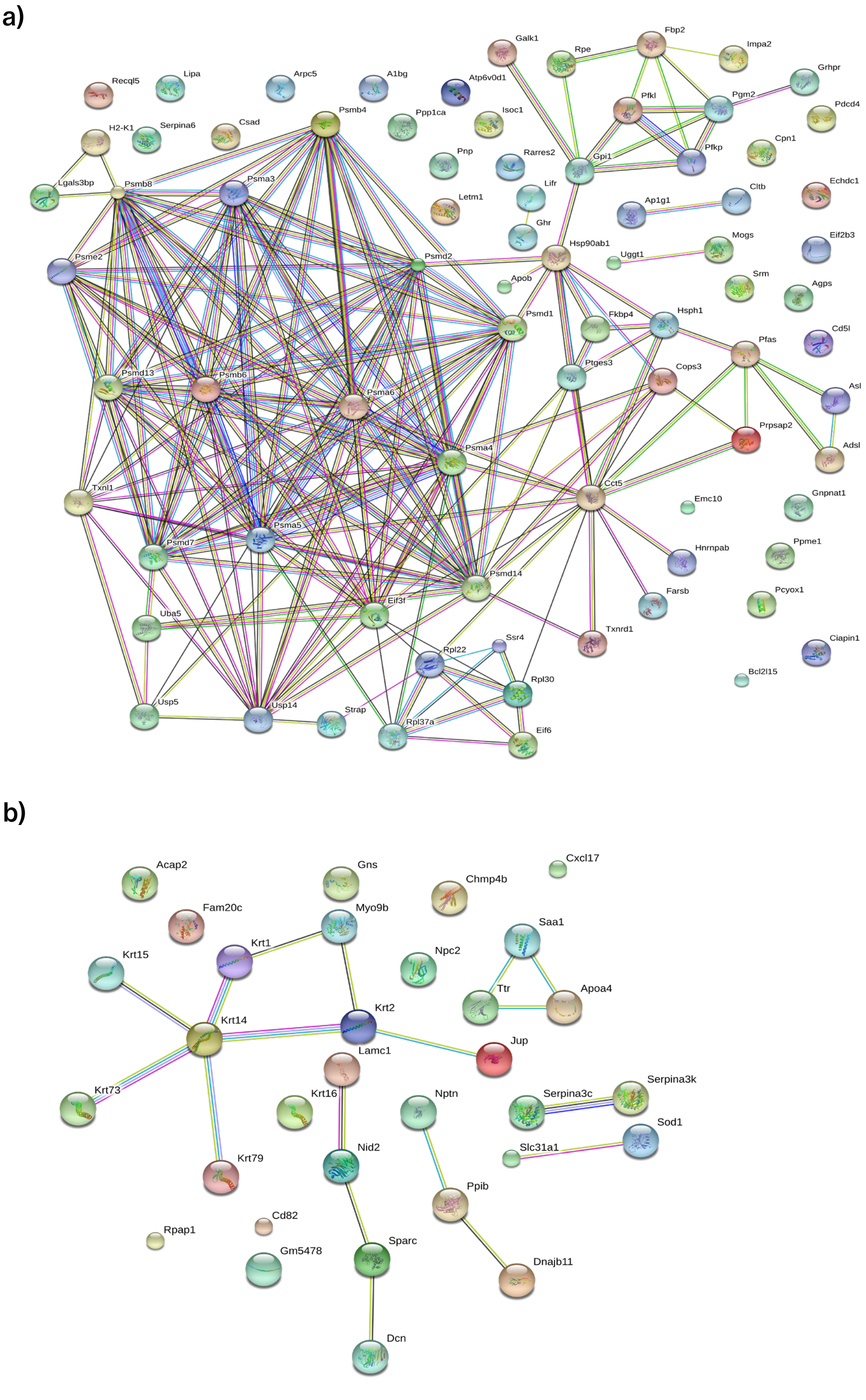


**Supplemental Figure 5. Analysis of breast milk proteins in NOD.IgA KO vs NOD dams**

STRING analysis of (a) significantly up-regulated and (b) significantly down-regulated proteins in NOD.IgA KO dams, where nodes indicate proteins and lines between nodes indicate known associations between those proteins.
